# Open Access Repository-Scale Propagated Nearest Neighbor Suspect Spectral Library for Untargeted Metabolomics

**DOI:** 10.1101/2022.05.15.490691

**Authors:** Wout Bittremieux, Nicole E. Avalon, Sydney P. Thomas, Sarvar A. Kakhkhorov, Alexander A. Aksenov, Paulo Wender P. Gomes, Christine M. Aceves, Andrés Mauricio Caraballo-Rodríguez, Julia M. Gauglitz, William H. Gerwick, Tao Huan, Alan K. Jarmusch, Rima F. Kaddurah-Daouk, Kyo Bin Kang, Hyun Woo Kim, Todor Kondić, Helena Mannochio-Russo, Michael J. Meehan, Alexey V. Melnik, Louis-Felix Nothias, Claire O’Donovan, Morgan Panitchpakdi, Daniel Petras, Robin Schmid, Emma L. Schymanski, Justin J. J. van der Hooft, Kelly C. Weldon, Heejung Yang, Shipei Xing, Jasmine Zemlin, Mingxun Wang, Pieter C. Dorrestein

## Abstract

Despite the increasing availability of tandem mass spectrometry (MS/MS) community spectral libraries for untargeted metabolomics over the past decade, the majority of acquired MS/MS spectra remain uninterpreted. To further aid in interpreting unannotated spectra, we created a nearest neighbor suspect spectral library, consisting of 87,916 annotated MS/MS spectra derived from hundreds of millions of public MS/MS spectra. Annotations were propagated based on structural relationships to reference molecules using MS/MS-based spectrum alignment. We demonstrate the broad relevance of the nearest neighbor suspect spectral library through representative examples of propagation-based annotation of acylcarnitines, bacterial and plant natural products, and drug metabolism. Our results also highlight how the library can help to better understand an Alzheimer’s brain phenotype. The nearest neighbor suspect spectral library is openly available through the GNPS platform to help investigators hypothesize candidate structures for unknown MS/MS spectra in untargeted metabolomics data.

## Introduction

When searching untargeted tandem mass spectrometry (MS/MS) metabolomics data using spectral libraries, on average only ∼5% of the data can be annotated (∼10% for human datasets). Unannotated spectra can arise due to incomplete coverage of the reference MS/MS spectral libraries of known compounds, including missing MS/MS spectra of different ion species, such as different ion forms, in-source fragments, and formation of multimers.^1–3^ We hypothesized that many of the unidentified ions originate from different but related known molecules. Those molecules could be a result of host or microbial metabolism or promiscuous enzymes that accept various analogous substrates during biosynthesis.^4^ To find related candidate ion species or to discover analogous MS/MS spectra from ions that originate from related molecules, strategies such as molecular networking^5^ and other analog searching strategies^3,6–10^ can be employed, for which molecular networking—a data visualization and interpretation strategy of MS/MS spectral alignment—in the Global Natural Products Social Molecular Networking (GNPS) environment^11^ is one of the most widely used tools.^12^

These strategies can also be used to generate new libraries of MS/MS reference spectra of potentially related MS/MS annotations from analog molecules that can subsequently be reused by the community. For example, small reference spectral libraries of human milk oligosaccharides^13^ and urine acylcarnitines^14^ were produced using an analog searching strategy (although the user licenses of these libraries restrict their redistribution). We hypothesized that the benefits of this approach could be further increased by considering analog matches across extremely large collections of MS/MS spectra to maximize the number of relevant spectrum links that can be found. Therefore, we have created a freely accessible and reusable MS/MS spectral library of MS/MS spectra related to identifiable molecules using molecular networking at the repository scale and created a nearest neighbor suspect spectral library to facilitate the annotation of mass spectrometry features that are present in public data.

## Results

### Nearest neighbor suspect spectral library creation

Using molecular networking, we have created a freely available and open-access mass spectral library of chemical analogs, referred to as the “nearest neighbor suspect spectral library.” The library was created from compatible public datasets deposited to GNPS/MassIVE,^11^ MetaboLights,^15^ and Metabolomics Workbench.^16^ In total, 521 million MS/MS spectra in 1335 public projects, with data from thousands of different organisms from diverse sources, including microbial culture collections, food, soil, dissolved organic matter, marine invertebrates, and humans, were used to compile the nearest neighbor suspect spectral library. Entries in this library, or “suspects,” were derived from unannotated spectra that were linked in a molecular network (based on spectral similarity) to an annotated spectrum by MS/MS spectral library searching and where the precursor ion mass difference between the two spectra was non-zero.

A hierarchical processing strategy was employed to compile the nearest neighbor suspect spectral library from repository-scale public MS/MS data (**Figure 1**). First, separate molecular networks were created for each dataset individually, while merging near-identical spectra and only keeping spectra that occur at least twice within the dataset to eliminate non-reproducible MS/MS spectra (**Figure 1, step 1**). Spectrum annotations were obtained at the individual dataset level by matching against 221,224 reference spectra available in the GNPS community spectral libraries (June 2021) using parameters consistent with a false discovery rate (FDR) <1%.^11^ The cosine similarity was calculated using filtered spectra (the precursor *m*/*z* peak was removed and only the top 6 most intense ions in every 50 *m*/*z* window were included), and spectrum matches with a cosine score of 0.8 or higher and a minimum of 6 matching ions were accepted. Second, a global molecular network was created from all of the individual networks using the GNPS modified cosine similarity (**Figure 1, step 2**). Finally, annotation propagations to the nearest neighbors were extracted from all molecular networks to create the library of nearest neighbor suspects (**Figure 1, step 3**). To maximize the quality of the suspect annotations, suspects with infrequent mass offsets that occur fewer than ten times were excluded, as these are considered to be less-reproducible mass differences (**Supplementary Figure 1**). Finally, a representative number of the annotation propagations were validated through expert manual inspection. The compilation of a global molecular network by co-networking thousands of datasets is an inherently more powerful strategy to discover relationships between MS/MS spectra, and thus between their corresponding molecules, than independent molecular networking within separate datasets, as moving to the repository scale makes it possible to discover patterns that cannot be detected from individual datasets in isolation.^17^ For example, if molecules are transformed during metabolism, the unmodified form might only be present as an endogenous molecule in the originating organism, such as plant or animal-based food products, with a modified variant due to metabolism present in samples from humans that consumed these foods (**Figure 1**).

**Figure 1.**
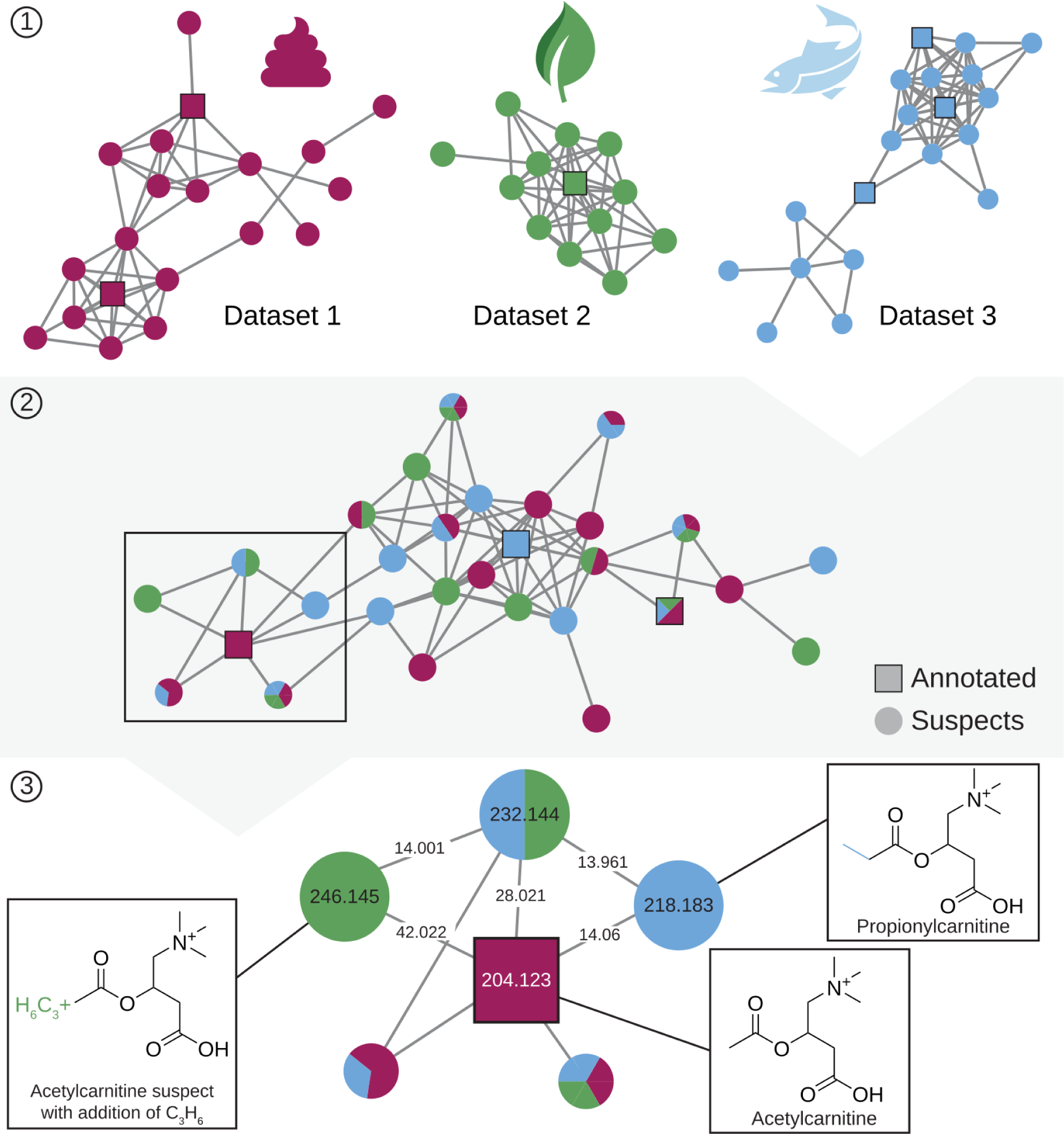
Creation of the nearest neighbor suspect spectral library. Overview of how the suspect library was created. Step 1: molecular networking of individual datasets. Step 2: co-networking of the 1335 datasets to create a global molecular network. Step 3: extract nearest neighbor suspects through annotation propagation to create the library.

In total, 87,916 unique MS/MS spectra and provenance to their matching analogs in the GNPS spectral libraries are included in the nearest neighbor suspect spectral library. Importantly, all of the nearest neighbor suspects are real spectra that occur in experimental data, whereas only a small portion (less than 10%) of reference MS/MS spectra in public and commercially available MS/MS spectral libraries have been observed in public data.^18^ To homogenize and extend the information available for the suspects, molecular formulas were determined using SIRIUS^19^ and BUDDY.^20^ The elemental composition of the suspects reflects the characteristics of known reference libraries (**Figure 2a**). For example, molecules that exclusively contain CH display poor ionization efficiency using electrospray ionization and are observed very rarely for both library types. Some suspects, such as common contaminants from sample vials, skin, or sodium formate clusters, as well as those related to endogenous molecules, such as fatty acids (e.g. vaccenic acid), bile acids (e.g. cholic acid), and lipids (e.g. phosphatidylcholines), are found in hundreds of public datasets and mass spectrometry files. In contrast, others, such as the natural products apratoxin, chelidonine, or marrubiin are observed less frequently (**Figure 2b, Supplementary Table 1**).

**Figure 2.**
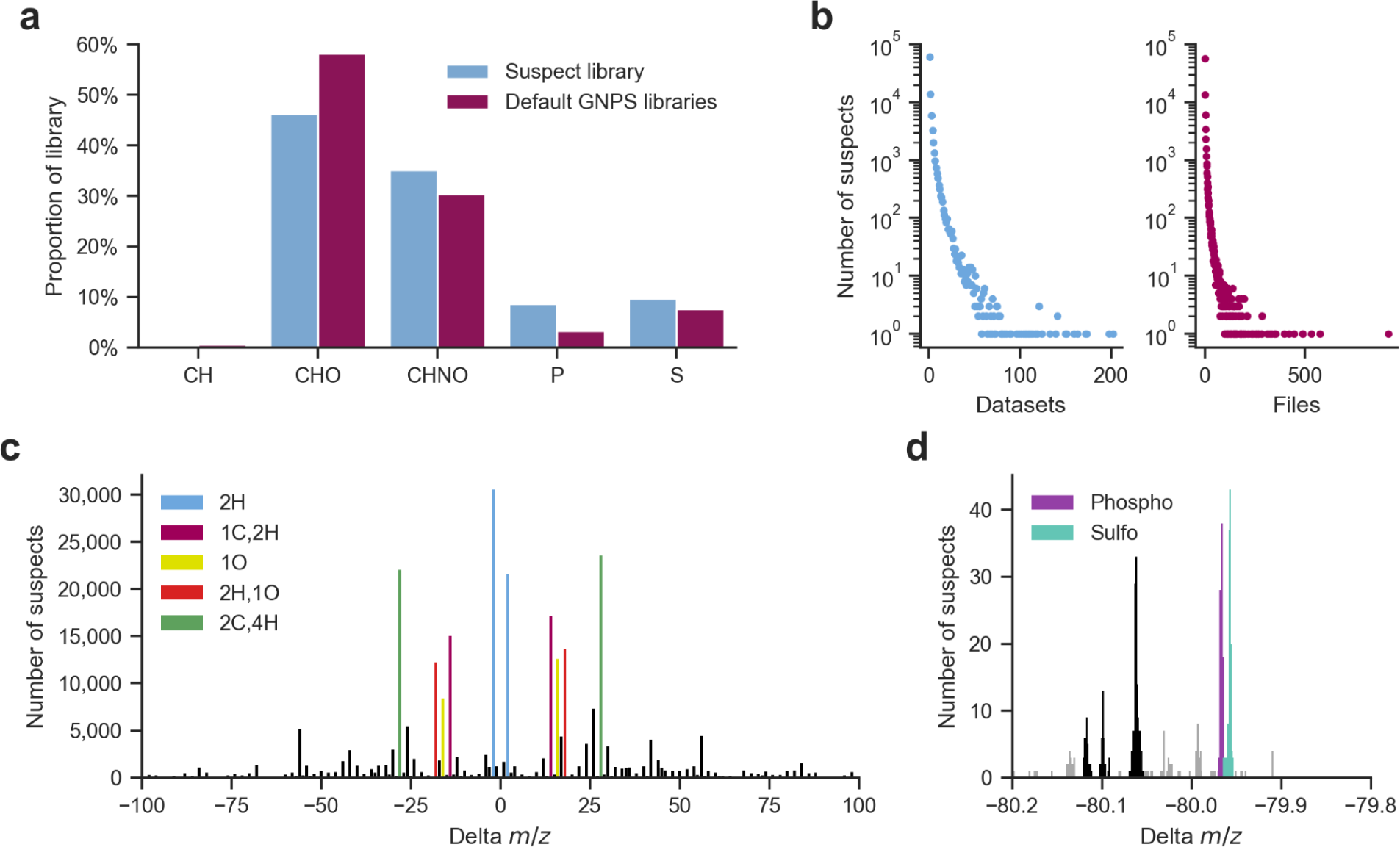
Composition of the nearest neighbor suspect spectral library. **a.** The composition of suspects that exclusively exist of CH, CHO, CHNO, or contain P or S compared to the reference libraries. **b.** Repeated occurrences of the suspects across datasets and files (i.e., individual LC-MS runs). **c.** Frequently observed mass offsets (delta masses between pairs of spectra) associated with the suspect library. **d.** Frequently observed mass offsets around a nominal mass of -80 Da.

There are 1350 frequent delta masses that occur in the nearest neighbor suspect spectral library (**Figure 2c, Supplementary Table 2**). When possible, the elemental composition of the delta masses and potential explanations, sourced from UNIMOD^21^—as many post-translational modifications or adducts that are observed in proteomics can also be found for small molecules—and a community-curated list of delta masses (**Supplementary Table 3**) are provided. The majority of delta mass explanations match the molecular formulas predicted by SIRIUS and BUDDY (**Supplementary Figure 1**), indicating the complementarity of these approaches to interpret the structural modifications that the suspects have undergone. The most common mass offsets observed in the suspect library correspond to a gain or loss of 2.016 Da, which can be interpreted as the gain or loss of 2H (e.g., a double bond or ring structure), followed by a gain or loss of 28.031 Da, 14.016 Da, 18.011 Da, and 15.995 Da, corresponding to C_2_H_4_ (e.g., di(de)methylation or (de)ethylation), CH_2_ (e.g., (de)methylation), H_2_O (e.g., water gain/loss), and O (e.g., (de)oxidation or (de)hydroxylation), respectively. However, 852 out of the 1350 mass offsets have not yet been explained (**Supplementary Figure 1**). For example, although these mass offsets occur less frequently, there are at least five repeatedly observed offsets with a nominal delta mass of -80 Da (**Figure 2d**), of which only phosphate loss (-79.966 Da) and sulfate loss (-79.957 Da) could currently be explained.

Spectral libraries are typically created by acquiring spectral data for pure standards, and reference MS/MS spectra have associated information on the precursor ions, compound names, and, when available, the molecular structures. In contrast, because the nearest neighbor suspect spectral library was compiled in a data-driven fashion, exact molecular structures are not known. Instead, the provenance of the suspect MS/MS spectra is described by their relationships to spectra that have an annotation, including the name and structure of the nearest neighbor MS/MS annotations and the observed pairwise delta masses. This is complemented by computed molecular formulas and the elemental composition and potential explanation of the delta masses, as determined by matching against a curated list of delta masses. Suspects thus represent unknown molecules that are likely structurally related to reference molecules annotated using spectral library searching, with the location of the structural modification generally unspecified. Without any additional information, this is in agreement with a level 3 annotation (family level match) according to the Metabolomics Standards Initiative guidelines.^22^

### Suspects provide structural hypotheses for observed molecules

The nearest neighbor suspect spectral library covers various classes of molecules arising from both primary and specialized metabolism, including lipids, flavonoids, and peptides. A fundamental understanding of organic chemistry, mass spectral fragmentation, and awareness of the information that mass spectrometry can or cannot provide is key to achieve the deepest possible structural insights from the suspect library. To demonstrate how the nearest neighbor annotations can be used to propagate structural information, we highlight examples of acylcarnitines, apratoxin natural products, drug metabolism, flavonoids, and polymers in greater detail. Note that we do not discuss the stereochemistry of the suspect examples, as this information generally cannot be determined using mass spectrometry.

The first example involves several acylcarnitines, a group of molecules that plays a key role in mammalian—including human—energy cycling (**Figure 3**).^23^ Hexanoylcarnitine, C6:0, is formed from the condensation of carnitine with hexanoic acid, a linear fatty acid with six carbons and zero double bonds (**Figure 3a**). Based on expert interpretation of the MS/MS spectra for suspects related to hexanoylcarnitine, we were able to derive structural hypotheses for several related acylcarnitines (**Figure 3b**). The first suspect example was initially annotated as a hexanoylcarnitine but with a loss of 2.016 Da. This indicates that this suspect is likely a hexenoylcarnitine derived from a C6:1 fatty acid. Thus, the six carbon fatty acid tail now has one double bond, but the location of the double bond and its configuration (*E* vs *Z*) cannot be determined. The second acylcarnitine suspect was annotated as a hexanoylcarnitine with a gain of 5.955 Da, which corresponds to a gain of one C along with the loss of six hydrogens. The only structure that can match the acyl side chain is a planar benzoyl ester. The third acylcarnitine suspect example showed an addition of 114.068 Da, representing a carnitine with an acyl side chain that has two oxygens and twelve carbons, as derived from the mass difference and characteristic neutral losses for carnitine conjugates of dicarboxylic acids (179.121 Da and 207.130 Da),^24^ which is consistent with dodecanedioylcarnitine. Additionally, we also found two suspects related to hexanoylcarnitine that include a characteristic 3-hydroxy fragment ion with a mass of 145.050 Da^25^ (**Figure 3c**). The first of these was initially annotated as hexanoylcarnitine but with a loss of 12.036 Da. Although close in value, based on accurate mass defects, the observed mass difference does not correspond to a loss of C (12.000 Da), but rather a combination that corresponds to the loss of C_2_H_4_ and gain of O. As the typical carnitine fragmentation pattern is conserved,^14^ we can determine that these changes occur in the fatty acid portion of the molecule, and thus, that this is likely a hydroxybutanoic acid carnitine derivative, leading to the final interpretation of 3-hydroxybutyrylcarnitine. The second 3-hydroxy suspect derived from hexanoylcarnitine showed an addition of 15.995 Da, representing a possible oxidation that can be localized based on the characteristic 3-hydroxy fragment,^25^ resulting in a spectrum annotation of 3-hydroxyhexanoylcarnitine. These two 3-hydroxy suspects also show excellent correspondence to commercial standards that were subsequently purchased and measured, confirming our structural assignments (**Figure 3c**).

**Figure 3.**
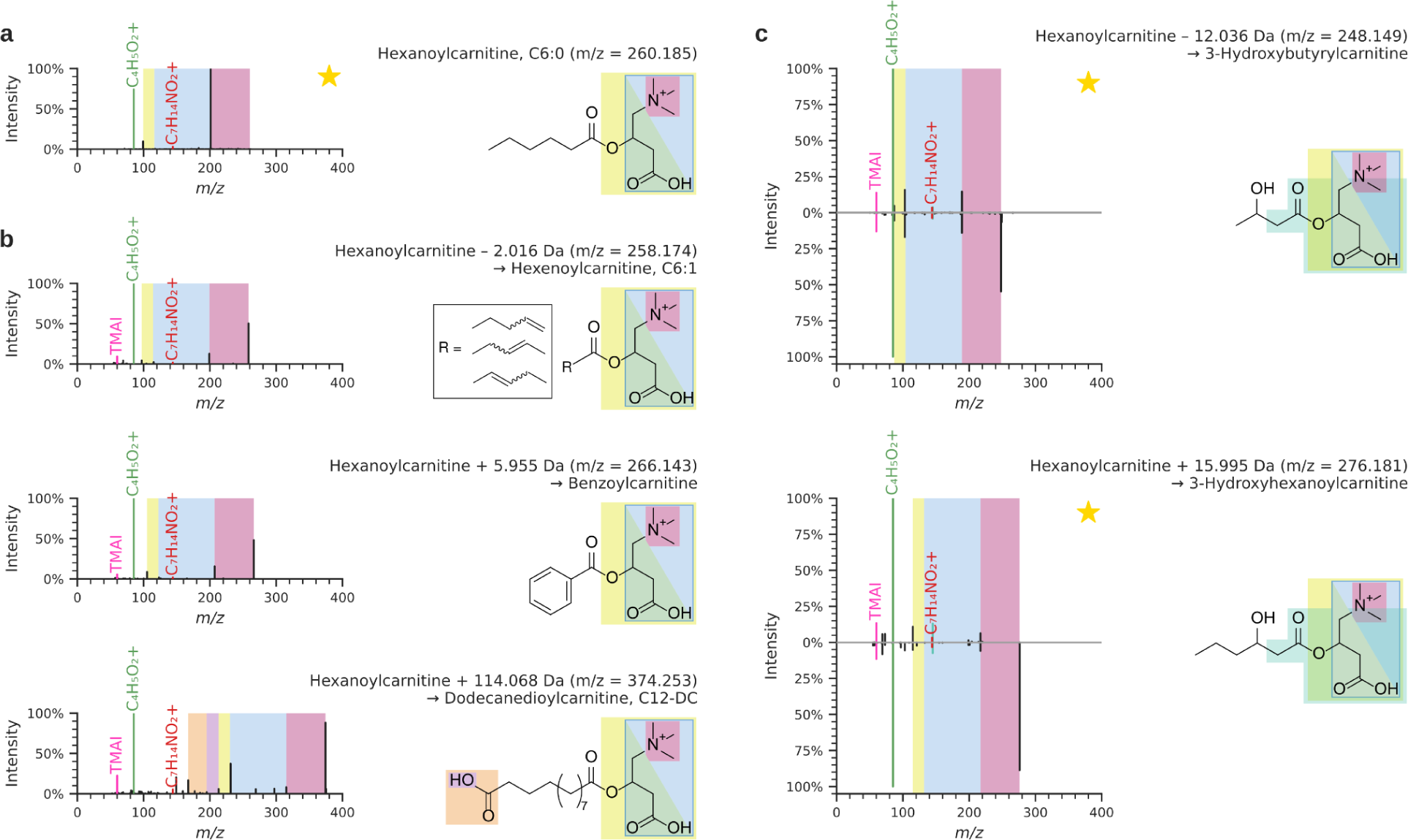
Novel acylcarnitine reference spectra obtained using the nearest neighbor suspect spectral library. Reference MS/MS spectra are indicated by ★. **a.** Reference MS/MS spectrum for hexanoylcarnitine originally included in the GNPS community spectral libraries. **b.** Nearest neighbor suspects related to hexanoylcarnitine. Annotations based on expert interpretation are: hexanoylcarnitine derived from a C6:1 fatty acid (unknown location of the double bond), benzoylcarnitine, and dodecanedioylcarnitine. **c.** Nearest neighbor suspects related to hexanoylcarnitine for 3-hydroxybutyrylcarnitine and 3-hydroxyhexanoylcarnitine (bottom) confirmed against commercial standards (top). The suspect MS/MS spectra show a very high cosine similarity of 0.9988 and 0.9927 to the reference MS/MS spectra for 3-hydroxybutyrylcarnitine and 3-hydroxyhexanoylcarnitine, respectively.

The suspect library is also informative for the analysis of more complex molecules. The apratoxin family of natural products was isolated from filamentous cyanobacteria, and has been investigated in a number of biological systems due to its potent antineoplastic activities.^26,27^ Using the suspect library to analyze a *Moorena bouillonii* cyanobacterial dataset achieved six additional spectrum annotations in the apratoxin molecular family (**Figure 4a**). A structural annotation can be determined for four of these based on comparisons to the MS/MS spectra of apratoxin standards. The four MS/MS spectra in **Figure 4b** show standards of purified apratoxin A, D, F, and C, while **Figure 4c** shows four apratoxin suspects with proposed structures. Some key substitutions observed are proline for *N*-methylalanine, methoxytyrosine for tyrosine, and dimethyl versus trimethyl polyketide initiating units. These substitutions are likely generated due to biosynthetic promiscuity commonly associated with multimodular hybrid non-ribosomal peptide synthetases-polyketide synthases.^28^ The apratoxin suspects that were observed are apratoxin A and F with loss of 30.010 Da, apratoxin A with loss of 26.015 Da, apratoxin A with loss of 28.031 Da, and apratoxin A and F with loss of 14.016 Da; corresponding to CH_2_O (e.g., methoxy loss), C_2_H_2_ or CH_2_ + C loss, C_2_H_4_ loss (e.g., dimethylation), and CH_2_ loss (e.g., methyl), respectively. The MS/MS spectra for four of the apratoxin suspects are shown in **Figure 4c**. Based on the fragmentation, the *m*/*z* difference corresponding to CH_2_ loss in apratoxin A is due to unmethylated tyrosine, which could be explained by inactivity of an *O*-methyl transferase during biosynthesis. The fragmentation for both apratoxins with the 30.010 Da loss supports that the *m*/*z* difference corresponding to methoxy loss is a result of phenylalanine incorporation by the associated adenylation domain rather than the methylated tyrosine observed in previously published apratoxin structures. Finally, although the loss of 26.015 Da is more complex, the other known apratoxins, together with their fragmentation, can be used to formulate a refined structural hypothesis. Compared to apratoxin A, the proline is likely substituted by an *N*-methylalanine, corresponding to a loss of C, and the trimethyl initiating unit is replaced by an isopropyl initiating unit. To obtain support for these modifications, isolation of this suspect (apratoxin A - 26.015 Da) was attempted from an extract of *Moorena bouillonii*; however, this resulted in a semi-pure fraction consisting of the suspect and small amounts of coeluting impurities. The semipure fraction was subjected to nuclear magnetic resonance (NMR) analysis (**Supplementary Figures 3–4**). Compared to NMR analysis of apratoxin A and consistent with the mass spectrometry interpretation, the NMR correlations associated with proline are lost and the NMR signals corresponding to *N*-methylation of alanine are now observed. Substructure analysis based on the MS/MS data revealed that the polyketide synthase portion of the apratoxin suspect differs by one methyl group. This is consistent with the suspect containing an isopropyl group, as observed in apratoxin C, rather than the *tert*-butyl group observed in nearly all of the other apratoxins.

**Figure 4.**
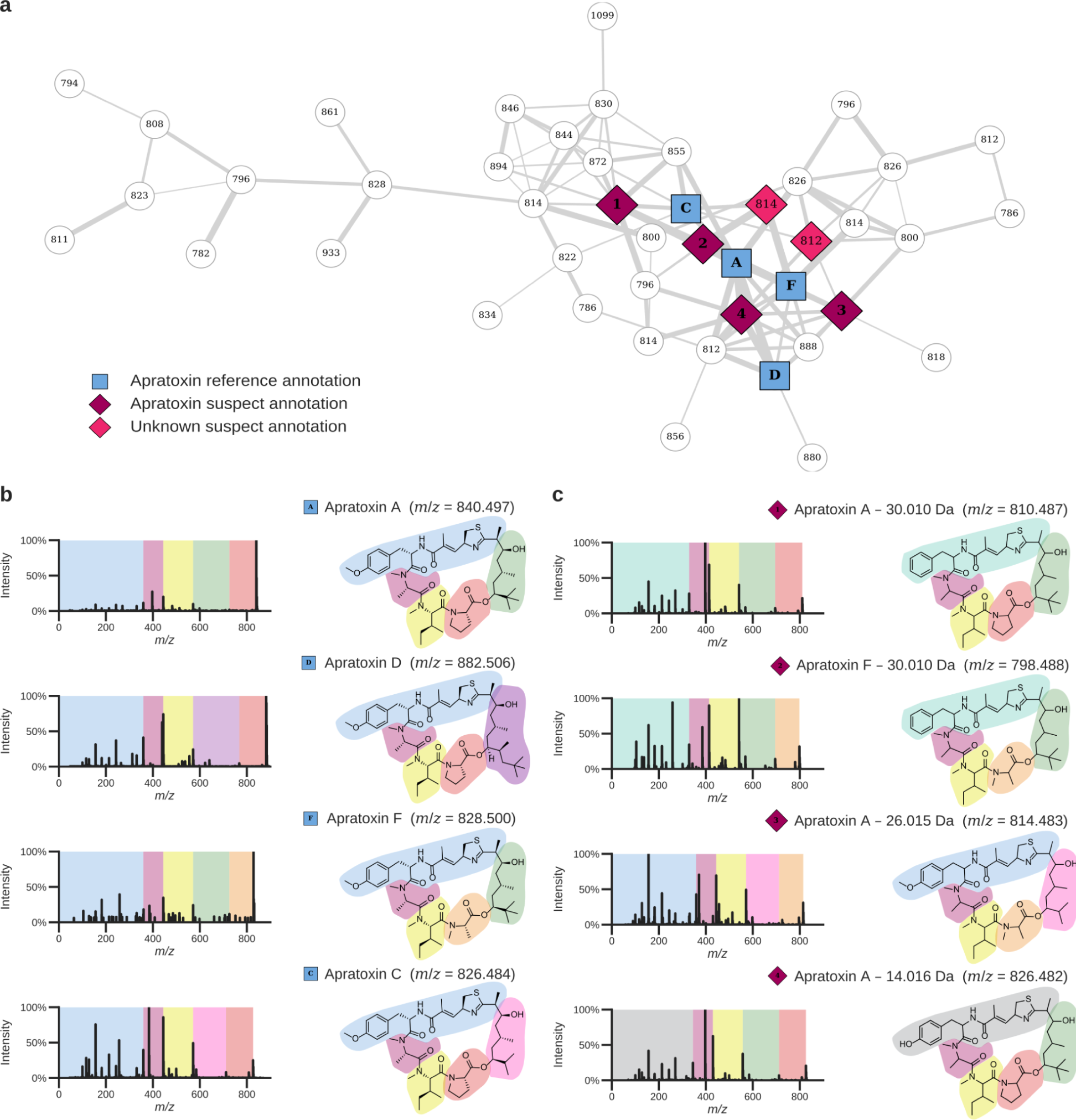
Novel apratoxin reference spectra obtained using the nearest neighbor suspect spectral library. **a.** Apratoxin cluster in a molecular network created from *Moorena bouillonii* crude extracts. The reference spectral library hits are shown by the blue squares (panel b). The purple and pink diamond nodes represent matches to the nearest neighbor suspect spectral library, with the purple diamonds matching the MS/MS spectra shown for which structures could be proposed (panel c). The white nodes are additional MS/MS spectra within the apratoxin molecular family that remained unannotated, even when including the suspect library. **b.** Reference MS/MS spectra and molecular structures of known apratoxins. **c.** MS/MS spectra and structural hypotheses for four novel apratoxin suspects. All four apratoxin suspects were derived from the tropical marine benthic filamentous cyanobacterium *Moorena bouillonii*, which is known to produce apratoxins (MSV000086109).

The suspect library also contains modified versions of known drugs that can arise due to in-source fragmentation, the formation of different ion species, incomplete synthesis or biosynthesis of the active ingredient that arises during manufacturing of the drug, or modifications introduced due to metabolism. An example is a suspect found in a human breast milk dataset matching the antibiotic azithromycin (**Supplementary Figure 5**).^29^ The suspect is 14.015 Da lighter, consistent with a CH_2_ loss. Based on inspection of the MS/MS data, it is possible to tentatively assign this loss of CH_2_ to the methoxy group in the cladinose sugar, based on the presence of a hydroxy loss and absence of a methoxy loss.

Next, MS/MS data from medicinal plants listed in the Korean Pharmacopeia were analyzed using molecular networking. Several flavonoid diglycosides containing pentoses and hexoses were detected using MS/MS spectral library searching, with the default GNPS libraries providing ten spectrum annotations in this molecular family and the suspect library contributing annotations for 27 diglycoside analogs (**Supplementary Figure 6**).^30^ Visual inspection of the MS/MS spectra indicated several modifications to formulate structural hypotheses for these suspects. For example, the apigenin-8-*C*-hexosylhexoside suspect with a delta mass of -30.019 Da corresponds to apigenin-8-*C*-pentosylhexoside. The presence of a pentose, instead of a hexose, is indeed consistent with the loss of CH_2_O.

Finally, analysis of closely related polymeric substances resulted in a substantial increase in annotations (**Supplementary Figure 7**). In an indoor chemistry environmental study,^31^ where a house was sampled before and after a month of human occupancy, there was a single spectrum match using the default GNPS libraries, to *p*-tert-octylphenol pentaglycol ether. Incorporating the suspect library added 55 matches that are related to polyethers, and that could be interpreted as part of a molecular family containing polymers. Thus, matching to the default GNPS spectral libraries alone gave the erroneous impression that there were only a few octylphenol-polyethylene glycol molecules detectable within the house, while the suspect library revealed that there is a large and diverse group of them.

In conclusion, these examples highlight how annotations provided by MS/MS spectral libraries, including the nearest neighbor suspect spectral library, can assist in providing structural hypotheses at the molecular family level for observed molecules.

### MS/MS spectrum annotation increases provide new biomedical insights

To evaluate the spectrum annotation performance of the nearest neighbor suspect spectral library, we performed spectral library searching of public untargeted metabolomics data on GNPS/MassIVE (**Figure 5a**). For the 1335 public datasets included during the creation of the suspect library, the default GNPS libraries resulted in an average MS/MS spectrum match rate of 5.5% (median 3.6%). Inclusion of the suspect library boosted the MS/MS spectrum match rate to 9.3% (median 6.4%), corresponding to 19 million additional spectrum matches. While these datasets were used to generate the suspect library, a similar increase in spectrum match rate was achieved for independent test data that were not part of the molecular networks from which the suspect library was compiled. For 72 datasets that were publicly deposited after the creation of the suspect library, the average spectrum match rate using the default GNPS libraries was 5.7% (median 4.7%), which increased to 8.9% (median 7.5%) when including the suspect library. Furthermore, we evaluated the performance of the suspect library for samples of different origins as recorded using the ReDU metadata system (**Figure 5b**).^17^ For 45,845 raw files from 179 datasets with controlled vocabularies for sample information, such as animal (including human), bacterial, fungal, environmental, food, and plant samples, the suspect library consistently achieved an increased spectrum match rate, ranging from a 1.7 ± 0.3 fold increase in interpreted spectra for food data to 3.0 ± 0.7 fold increase for environmental samples (mean and standard deviation).

**Figure 5.**
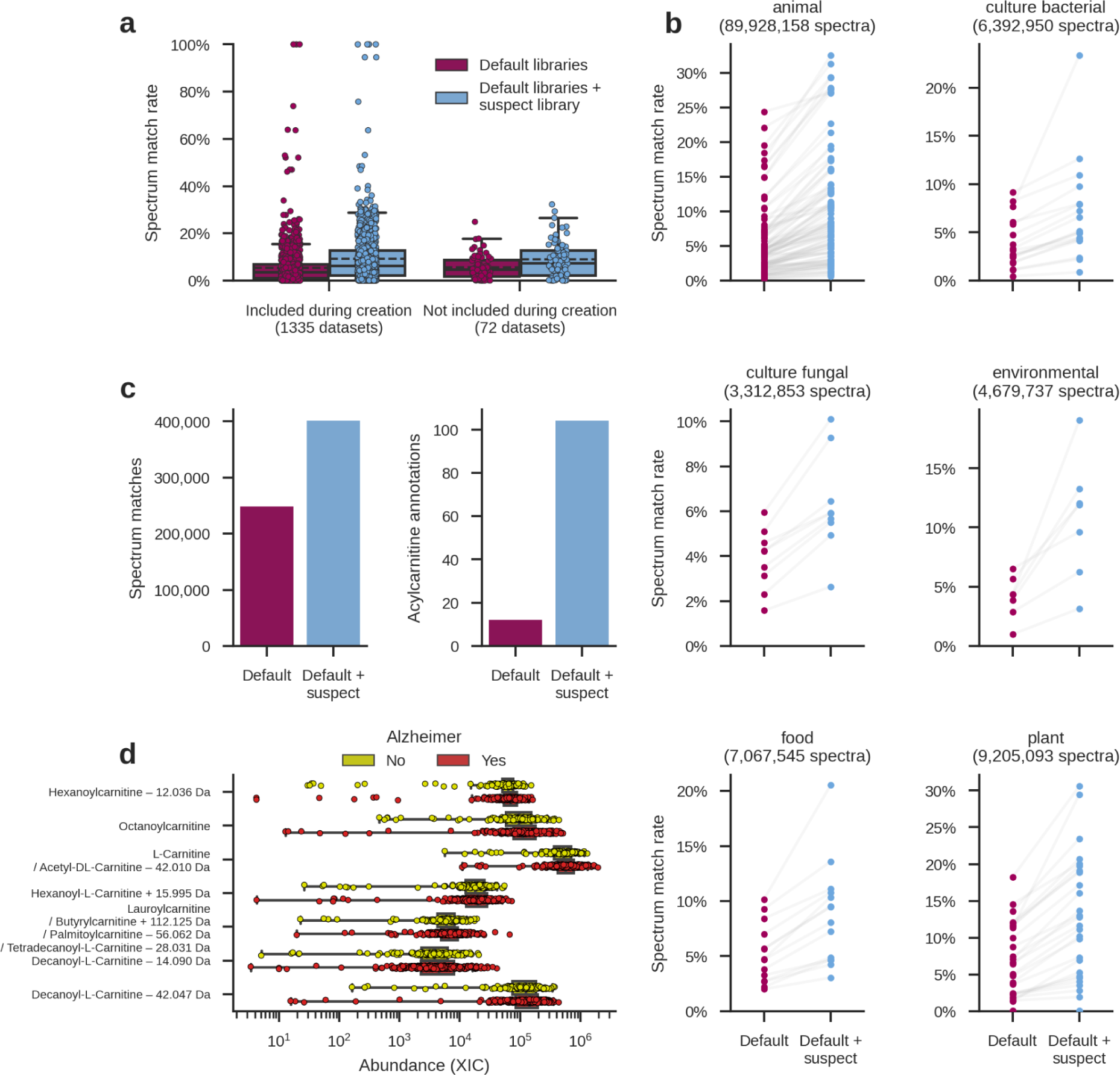
Impact of the nearest neighbor suspect spectral library on spectrum matches to enable the formulation of structural hypotheses. **a.** The MS/MS spectrum match rate with and without the suspect library for 1407 public datasets on GNPS/MassIVE. The full center line indicates the median values, and the dashed center line indicates the mean values. The box limits indicate the first and third quartiles of the data, and the whiskers extend to 1.5 times the interquartile range. **b.** The MS/MS spectrum match rate for different types of datasets with and without the suspect library. The data comes from 45,845 raw files in 179 datasets with known sample types recorded using the ReDU metadata system.^17^ **c.** MS/MS matches to an untargeted metabolomics human brain dataset from Alzheimer’s disease patients (n=360) and healthy subjects (n=154) with and without the suspect library. **d.** Differentially abundant carnitines for Alzheimer’s disease patients (Benjamini-Hochberg corrected p-value < 0.05). The suspect library was able to identify four additional mass spectrometry features as acylcarnitines, which would have remained unannotated matched only against the default GNPS libraries. The box limits indicate the first and third quartiles of the data, and the whiskers extend to 1.5 times the interquartile range.

To further demonstrate the utility of the suspect library, we focused on untargeted metabolomics data from 514 human brains with and without Alzheimer’s disease.^32^ Using the default GNPS libraries there were 248,317 MS/MS spectral library matches, corresponding to 1305 unique molecule annotations. Including the suspect library increased the number of spectrum matches to 401,039, covering 5184 unique molecule annotations (**Figure 5c**). One specific class of molecules that saw a particularly large increase in the number of annotations in this cohort was the acylcarnitines. There were 942 spectrum matches to 12 unique molecule annotations before the suspect library was included, but 1896 spectrum matches to 104 unique molecule annotations after inclusion of the suspect library.

We observed significant abundance differences for six acylcarnitines, as well as carnitine, when comparing brain metabolites between groups with and without Alzheimer’s diagnosis (**Figure 5d**). Three of those—carnitine, octanoylcarnitine, and lauroylcarnitine—could be annotated using the default GNPS libraries, while the remaining four metabolites could only be identified as acylcarnitines using the suspect library. The annotations of these seven metabolites were covered by three spectral library matches and eight suspect matches. When multiple spectra matched against the same metabolite, these annotations reinforced each other. For example, the carnitine annotation co-occurs with a match to the acetylcarnitine suspect with a loss of 42.010 Da. In this case, 42.010 Da corresponds to the mass of acetylation, which is lost in the suspect annotation, and therefore the suspect MS/MS spectrum represents carnitine itself. The four suspects that would otherwise remain unassigned as potential acylcarnitines are hexanoylcarnitine with loss of 12.036 Da (3-hydroxybutyrylcarnitine, **Figure 3c**), hexanoylcarnitine with addition of 15.995 Da (3-hydroxyhexanoylcarnitine, **Figure 3c**), decanoyl-L-carnitine with loss of 14.090 Da (-3C,-10H,+2O in the acyl chain), and decanoyl-L-carnitine with loss of 42.047 Da (-3C,-6H in the acyl chain). The first two suspects are related 3-hydroxy acylcarnitines that have 3-hydroxy-butyrate and 3-hydroxy-hexonate as the acyl side chain, whose predictions matched to data from commercial standards (**Figure 3c**). The other two suspects are consistent with DC7:1 and C7:0 acylcarnitines.^33^ These observations provide additional support that there are different fatty acids—now also including 3-hydroxy and odd-chain fatty acids—that are transported as carnitine derivatives in Alzheimer’s disease brains in comparison to healthy brains.^34,35^

## Discussion

The annotation of untargeted metabolomics data is based on reference spectral libraries. However, because many known compounds and previously undiscovered analogs of compounds are unavailable as reference standards, alternative approaches are required to interpret such MS/MS spectra. Here we have introduced a data-driven approach to compile an extensive nearest neighbor suspect spectral library. This library consists of 87,916 unique MS/MS spectra and can be freely downloaded as a Mascot generic format (MGF) file from the GNPS website. Additionally, through its direct integration in the spectral library searching and molecular networking functionality on the GNPS platform, the scientific community can incorporate the nearest neighbor suspect spectral library in their data analyses to formulate structural hypotheses.

The nearest neighbor suspect spectral library is closely related to analog searching. As suspects are derived from unannotated spectra that were linked in a molecular network to an annotated spectrum by MS/MS spectral library searching, conceptually analog searching can be used to directly annotate the MS/MS spectra as well. However, the nearest neighbor suspect spectral library has several ways it complements and/or brings advantages to analog searching.

First, analog searching is computationally very expensive due to the massive search space that needs to be considered by opening up the precursor mass tolerance. As such, optimized algorithms^36,37^ or even specialized hardware^38^ are required to be able to do this efficiently. In contrast, the nearest neighbor suspect spectral library seamlessly works with standard spectral library searching procedures that are ubiquitously available. Second, analog searching suffers from an increased rate of false positive annotations, as random high-scoring matches are more likely to occur when considering a very large search space. In contrast, we explicitly safeguarded the quality of the nearest neighbor suspect spectral library by using stringent approaches towards spectrum matching and data filtering. Third, it can be challenging to interpret individual MS/MS spectrum annotations from analog searching, which generally involves a manual process. The nearest neighbor suspect spectral library addresses this by integrating relevant information from various sources and tools, including predicted molecular formulas and delta mass interpretations, to facilitate its annotations. In essence, this library contextualizes MS/MS spectrum annotations obtained from the nearest neighbor suspect spectral library within the full library and even the global molecular network from which it was compiled, making this an inherently more efficient strategy than analog searching.

Entries in the nearest neighbor suspect spectral library are not obtained by measuring pure reference standards. Therefore, it is important to consider that, initially, the exact molecular structure of the suspects is undetermined. Nevertheless, the suspect library includes essential information that can help to interpret MS/MS data that would otherwise remain entirely unexplored. Additionally, all of the spectra that are part of the suspect library have been detected experimentally and occur in biological data. In contrast, only a minority of the compounds contained in reference spectral libraries are actually observed in public data, indicating a mismatch between the laborious reference library creation efforts and the practical needs of metabolomics researchers. Consequently, incorporating the nearest neighbor suspect spectral library significantly increases the spectrum match rate across a wide variety of sample types. We have demonstrated how careful investigation of the suspects can provide highly detailed interpretations, and we anticipate that similar community contributions will be used to add and confirm further suspect annotations. Finally, when future studies uncover biologically relevant suspects, their molecular identities, including the location of modifications and stereochemical features, might be refined by measuring orthogonal properties, such as collision cross-section by ion mobility spectrometry or using genome mining, when possible. Ultimately, as is the case for all spectrum annotations, experimental validation of the complete molecular stereostructure requires either a reference standard or further isolation followed by structure elucidation by NMR, X-ray crystallography, or cryogenic electron microscopy experiments.

## Acknowledgements

This research was supported in part by BBSRC-NSF award 2152526. This research was supported in part by National Institutes of Health awards R01 GM107550, U19 AG063744, U01AG061359, R03 CA211211, P41 GM103484, T32 HD123456. This research was supported in part by the National Institute of Aging’s Accelerating Medicines Partnership for AD (AMP-AD) and was supported by NIH grants 1R01AG069901-01A1, U01AG061357, as well as by the Alzheimer Gut Microbiome Project grant 1U19AG063744. This research was supported in part by federal award DE-SC0021340 subaward 1070261-436503. This research was supported in part by the Gordon and Betty Moore Foundation through grant GBMF7622. This research was supported in part by the Intramural Research Program of National Institute of Environmental Health Sciences of the National Institutes of Health (ZIC ES103363). This research was supported in part by the National Center for Complementary and Integrative Health of the NIH under award number F32AT011475 to NEA. ELS & TK acknowledge funding support from the Luxembourg National Research Fund (FNR) for project A18/BM/12341006. MW was partially supported by the US Department of Energy Joint Genome Institute operated under Contract No. DE-AC02-05CH11231. DP was supported by the Deutsche Forschungsgemeinschaft (DFG) through the CMFI Cluster of Excellence (EXC 2124). SAK was supported by the Fund for Financing Science and Supporting Innovation under the Ministry of Innovative Development of the Republic of Uzbekistan. KBK was supported by the National Research Foundation of Korea (NRF) grants funded by the Ministry of Science and ICT (NRF-2020R1C1C1004046). HM-R acknowledges the Brazilian National Council for Scientific and Technological Development (CNPq, #142014/2018-4) and the Brazilian Fulbright Commission for the scholarships provided. LFN has been supported by the French government, through the UCA^J.E.D.I.^ Investments in the Future project managed by the National Research Agency (ANR) with the reference number ANR-15-IDEX-01. JJJvdH was supported by an ASDI eScience grant from the Netherlands eScience Center (ASDI.2017.030). COD was supported by EMBL core funds. The Alzheimer’s disease metabolomics data was funded wholly or in part by the Alzheimer’s Gut Microbiome Project (AGMP) NIH grant U19AG063744 awarded to Dr. Kaddurah-Daouk at Duke University in partnership with a large number of academic institutions. More information about the project and the institutions involved can be found at https://alzheimergut.org/meet-the-team/.

## Author contributions statement

PCD conceptualized and supervised the work. COD and CMA helped transfer and convert data from MetaboLights. WB, MW, and PCD created the methodology to compile the nearest neighbor suspect spectral library from molecular networking data. WB, JMG, and MW developed the software. WB, NEA, SPT, SAK, AAA, PWPG, AMCR, JMG, AKJ, TK, HM-R, MJM, LFN, MP, DP, RS, RS, ELS, and JJJvdH validated entries in the suspect spectral library and evaluated its identification performance. NEA, SPT, SAK, AAA, and PWPG provided case studies to demonstrate the utility of the tool. MW provided computational resources. CMA, CO, MP, and JZ performed data curation. AMCR acquired the acylcarnitine standards MS/MS data. WHG supervised acquisition of the *Moorena bouillonii* MS/MS data. KBK, HWK, and HY acquired the medicinal plants from the Korean Pharmacopeia MS/MS data. AAA and AM acquired the HomeCHEM MS/MS data. RFKD supervised acquisition of the ROSMAP metabolomics data and links to ADMC and AMP-AD consortia. MJM, MP, KCW, and JZ processed and prepared the ROSMAP samples and acquired the MS/MS data. WB, NEA, SPT, AAA, PWPG, CMA, MJM, and PCD wrote the manuscript. All authors reviewed and edited the manuscript.

## Methods

### Integration of Metabolights into GNPS/MassIVE

As a joint effort of the European Bioinformatics Institute (EMBL-EBI) and the GNPS/MassIVE teams, approximately 10,000 LC-MS/MS samples acquired in positive ion mode were imported from MetaboLights^15^ into the GNPS/MassIVE repository by mirroring relevant files from MetaboLights on GNPS/MassIVE. These files represent over a hundred studies containing data from biologically diverse backgrounds, including but not limited to human, fungus, various bacterial and microbial species, and ecological samples. The data consist of both metabolomics and lipidomics samples.

### GNPS living data molecular networking

The nearest neighbor suspect spectral library was derived from molecular networking results as performed by GNPS’s “living data” functionality, which periodically reanalyzes all publicly available untargeted metabolomics data on GNPS/MassIVE.^11^ The living data analysis (update performed on November 17, 2020) includes results for 1335 datasets, corresponding to 520,823,130 million MS/MS spectra. Spectrum clustering grouped 168,193,526 MS/MS spectra in 8,543,020 clusters while discarding 352,629,604 singleton spectra, of which 454,091 cluster representatives could be annotated using spectral library searching against the default GNPS spectral libraries (5.3% annotation rate), and which formed a molecular network consisting of 13,179,147 spectrum pair edges.

This analysis consisted of two phases of spectrum clustering using MS-Cluster^39^ and molecular networking. First, spectra were networked within each individual dataset. Per-dataset molecular networking outputs are available on the MassIVE repository with dataset identifier <u>MSV000084314</u>. Next, a second round of molecular networking was performed on the combined consensus spectra for all datasets generated from the first molecular network.

Spectra were preprocessed by removing all MS/MS fragment ions within +/- 17 Da of the precursor *m*/*z.* Only the top 6 most abundant ions in every 50 *m*/*z* window were retained. The first round of molecular networking used a precursor mass tolerance of 2.0 *m*/*z*, a fragment mass tolerance of 0.5 *m*/*z*, and three rounds of MS-Cluster clustering with mixture probability threshold 0.05. Thresholds for the second round of molecular networking were modified due to computational and memory constraints, and consisted of a precursor mass tolerance of 0.1 *m*/*z* and a fragment mass tolerance of 0.1 *m*/*z*. MS-Cluster used the standard cosine similarity to group near-identical MS/MS spectra, whereas the molecular networking analyses employed a GNPS modified cosine similarity that takes directly matching ions into account as well as ions that are shifted according to the precursor mass difference.^9^ The molecular networking used a minimum cosine similarity of 0.8, minimum six matched peaks, only considered clusters that consist of at least two MS/MS spectra, and retained the ten strongest edges for each node in the molecular network.

Spectral annotations were obtained through spectral library searching against the default GNPS spectral libraries (GNPS Collections Bile Acid Library 2019,^11^ CASMI,^40^ Dereplicator Identified MS/MS Spectra,^41^ GNPS Collections Miscellaneous,^11^ Pesticides, EMBL Metabolomics Core Facility,^42^ Faulkner Legacy Library provided by Sirenas MD, GNPS Library,^11^ NIH Clinical Collection 1, NIH Clinical Collection 2, NIH Natural Products Library Round 1,^43^ NIH Natural Products Library Round 2,^43^ Pharmacologically Active Compounds in the NIH Small Molecule Repository, GNPS Matches to NIST14,^11^ PhytoChemical Library, FDA Library Pt 1, FDA Library Pt 2, HMDB,^44^ LDB Lichen Database,^45^ Massbank Spectral Library,^46^ Massbank EU Spectral Library, MIADB Spectral Library,^47^ Medicines for Malaria Venture Pathogen Box, Massbank NA Spectral Library, Pacific Northwest National Lab Lipids,^48^ ReSpect Spectral Library,^49^ Sumner/Bruker), which contained 221,224 reference MS/MS spectra (June 2021). Settings for the living data spectral library searching step included a precursor ion tolerance of 2.0 *m*/*z*, a fragment ion tolerance of 0.5 *m*/*z*, a minimum cosine similarity of 0.7, and minimum six matched peaks.

### Nearest neighbor suspect spectral library creation

High-quality MS/MS spectra were extracted from the GNPS living data molecular network to compile the nearest neighbor suspect spectral library. Suspects were derived from spectrum pairs for which only one of the spectra was identified during spectral library searching and both spectra have a non-zero precursor mass difference. In this case, the unidentified spectrum was included in the nearest neighbor suspect spectral library, as it corresponds to a previously unknown molecule that is structurally related to the reference molecule identified using spectral library searching. Strict filtering thresholds were used to avoid inclusion of incorrect entries: spectrum–spectrum matches required a maximum precursor mass tolerance of 20 ppm, a minimum cosine similarity of 0.8, and minimum six matched ions.

To homogenize and extend the information available for the suspects, their molecular formulas were determined using SIRIUS^19^ and BUDDY.^20^ For the SIRIUS (version 4.5.2) analysis, only MS/MS data were used as input and the precursor mass tolerance was set to 10 ppm for Orbitrap spectra and 25 ppm for Q-TOF spectra. For the BUDDY (version 1.3) analysis, a precursor mass tolerance of 5 ppm for FT-ICR spectra, 10 ppm for Orbitrap spectra, and 25 ppm for Q-TOF spectra was used. The fragment mass tolerance was set to twice the precursor mass tolerance and no database restriction was applied. All other settings were kept at their default values. If BUDDY could not annotate MS/MS spectra with molecular formulas while only considering CHNOPS elements, the spectra were subsequently reannotated while considering additional elements: CHNOPSFClBrI. Molecular formulas predicted by SIRIUS and BUDDY agreed for 32,302 suspects, while SIRIUS predicted a different molecular formula for 32,508 suspects and BUDDY for 47,508 suspects. In case SIRIUS and BUDDY predicted different molecular formulas, both were included in the suspect name (see below).

Additionally, the observed precursor mass differences were calibrated and matched to putative modification explanations contained in the UNIMOD database^21^ and a manually compiled list of modifications and their mass differences (**Supplementary Table 3**). Suspects whose delta mass occurred fewer than ten times were discarded, as true modifications are expected to occur repeatedly for different molecules, while suspects with infrequent mass differences more likely correspond to spurious matches.

To ensure that the provenance of the suspects to the matched reference molecules compared to which they are annotated based on spectral similarity is properly understood, their names are of the form: “Suspect related to [*compound name*] (predicted molecular formula: [*molecular formula SIRIUS and/or BUDDY*]) with delta *m*/*z* [*positive (addition) or negative (loss) delta m/z*] (putative explanation: [*modification*]).” In case multiple propagations to different reference spectra are available, information for all matches is included.

### Spectrum annotation using the nearest neighbor suspect spectral library

The spectrum annotation performance of the nearest neighbor suspect spectral library was assessed by large-scale spectral library searching using the default GNPS spectral libraries excluding and including the nearest neighbor suspect spectral library on 1407 public datasets available on GNPS/MassIVE, consisting of a combined 592 million MS/MS spectra. Of these datasets, 1335 datasets were also included in the GNPS living data analysis from which the nearest neighbor suspect spectral library was compiled (521 million MS/MS spectra; see above) and 72 datasets were deposited at a later date and can be considered a completely independent test set (72 million MS/MS spectra). All searches used a precursor mass tolerance of 2.0 *m*/*z*, a fragment mass tolerance of 0.5 *m*/*z*, a minimum cosine similarity of 0.8, and minimum 6 matched peaks. Other options were kept at their default values.

### Evaluation of acylcarnitine suspects

#### Mass spectrometry analysis

Structural hypotheses for several suspect acylcarnitines were confirmed using reference standards based on spectral matches, accurate masses, and retention times: [(3R)-3-Hydroxybutyryl]-L-carnitine (Catalog No. 918639-76-6, Sigma-Aldrich Inc.), [(3R)-3-Hydroxyhexanoyl]-L-carnitine (Catalog No. 1469900-93-3, Sigma-Aldrich Inc.) and Hexanoyl-L-carnitine (Catalog No. 22671-29-0, Sigma-Aldrich Inc.). Standards were prepared at 1uM concentration. Untargeted LC-MS/MS acquisition was performed on a Vanquish Ultrahigh Performance Liquid Chromatography (UHPLC) system coupled to a Q-Exactive Hybrid Quadrupole-Orbitrap (Thermo Fisher Scientific, Bremen, Germany). Chromatographic separation was performed on a Kinetex 1.7 μm 100 Å pore size C18 reversed phase UHPLC column 50 × 2.1 mm (Phenomenex, Torrance, CA) with a constant flow rate of 0.5 mL/min. The following solvents were used during the LC-MS/MS acquisition: water with 0.1% formic acid (v/v), Optim LC/MS grade, Thermo Scientific (solvent A) and acetonitrile with 0.1% formic acid (v/v), Optima LC/MS grade, Thermo Scientific (solvent B). After injection of 1uL of sample into the LC system and eluted with isocratic gradient of 5% B from 0 to 1 mins and linear gradient from 5 to 100% B (1–7 min), 100% B (7–7.5 min), 100 to 5% B (7.5–8 min), 5% B (8–10 min). Data dependent acquisition mode was used for acquisition of MS/MS data with default charge state of 1. An inclusion list containing the following ions was used: *m/z* 260.18563 (molecular formula: C13H25NO4, start: 2.00 min, end: 3.00 min), *m/z* 248.14925 (molecular formula: C11H21NO5, start: 0.00 min, end: 1.00 min), *m/z* 276.18055 (molecular formula: C13H25NO5, start: 0.50 min, end: 1.50 min). Full MS was acquired using 1 microscan at a resolution of 35,000 at 200 *m*/*z*, automatic gain control (AGC) target 5e5, maximum injection time of 100 ms, scan range 100–1500 *m*/*z* and data acquired in profile mode. DDA of MS/MS was acquired using 1 microscan at a resolution of 35,000 at 200 *m*/*z*, AGC target 5e5, top 5 ions selected for MS/MS with isolation window of 2.0 *m/z* with scan range 200–2000 *m/z*, fixed first mass of 50 *m*/*z* and stepped normalized collision energy of 20, 30, and 40 eV, minimum AGC target 5e3, intensity threshold 5e4, apex trigger 2 to 15 s, all multiple charges included, isotopes were excluded, and a dynamic exclusion window of 10 s.

### Evaluation of apratoxin suspects

#### Mass spectrometry analysis

Apratoxin suspects were investigated in the context of *Moorena bouillonii*, a tropical marine benthic filamentous cyanobacterium. The mass spectrometry data were derived from both field-collected and laboratory-cultured biomass of *Moorena bouillonii* (MassIVE dataset identifier MSV000086109). A number of collections are represented in this dataset, including those originating from sites around Guam, Saipan (Commonwealth of the Northern Mariana Islands), Palmyra Atoll, Papua New Guinea, American Samoa, Kavaratti (Lakshadweep, India), the Paracel Islands (Xisha, China), the Solomon Islands, and the Red Sea (Egypt). The biomass from each of the samples was extracted using 2:1 dichloromethane and methanol. The crude extracts were concentrated and resuspended in acetonitrile, followed by a desalting protocol using C18 SPE with acetonitrile. Samples were then resuspended in methanol containing 2 μM sulfamethazine as an internal standard. Untargeted metabolomics was performed using an UltiMate 3000 liquid chromatography system (Thermo Scientific) coupled to a Maxis Q-TOF (Bruker Daltonics) mass spectrometer with a Kinetex C18 column (Phenomenex). Data were collected in positive ion mode using data-dependent acquisition. All solvents used were LC-MS grade.

#### Molecular networking

Molecular networking and spectral library searching were performed using the GNPS platform as described above. Settings included a precursor mass tolerance of 2.0 *m*/*z*, fragment mass tolerance of 0.5 *m*/*z*, minimum cosine similarity of 0.7, and minimum 6 matched peaks. Data visualization was performed using the Metabolomics USI interface^50^ and spectrum–spectrum matches were evaluated manually to develop hypotheses regarding the structure of apratoxin analogs that were annotated using the nearest neighbor suspect spectral library.

#### Cyanobacterial culture

*Moorena bouillonii* PNG5-198 was initially collected by scuba in 3–10 m of water off the coast of Pigeon Island, Papua New Guinea (S4 16.063’ E152 20.266’) in May 2005. Live cultures have been maintained in SWBG-11 media under laboratory conditions at 27°C and a 16/8 h light/dark schedule. Biomass for *Moorena bouillonii* was obtained through ongoing laboratory culture.

#### Extraction and isolation of apratoxins

The cultured biomass was extracted using 2:1 CH_2_Cl_2_/MeOH affording 241.4 mg of organic extract. The extract was then subjected to vacuum liquid chromatography (VLC) on silica gel (type H, 10–40 μm) using normal phase solvents in a stepwise gradient of hexanes/EtOAc and EtOAc/MeOH, resulting in nine fractions (A-I). The fraction eluting with 25% MeOH/75% EtOAc (fraction H) had a mass of 21.6 mg. This fraction was found to have the characteristic MS/MS signatures of the apratoxins and was selected for further purification using reversed-phase HPLC. A Phenomenex Kinetex C18 5*μ*m 100Å 100 × 4.6 mm column with a 3 mL/min was used to obtain 1.2 mg of semipure suspect (apratoxin A - 26.015 Da) and 2.1 mg of semipure apratoxin A.

#### NMR spectroscopy

1H NMR and 2D NMR spectra were obtained on a Bruker Advance III DRX-600 NMR with a 1.7 mm dual tune TCI cryoprobe (600 MHz and 150 MHz for ^1^H and ^13^ C, respectively). NMR spectra were referenced to residual solvent CDCl_3_ signals as an internal standard. NMR spectra were processed using MestReNova (Mnova 14.2.3, Mestrelab Research).

### Evaluation of azithromycin suspects

#### Mass spectrometry analysis

The presence of azithromycin suspects was investigated using human breast milk data (MassIVE dataset identifier MSV000081432).^29^ Human milk samples were extracted using 80:20 methanol and water. Untargeted metabolomics was performed using an UltiMate 3000 liquid chromatography system (Thermo Scientific) coupled to a Maxis Q-TOF (Bruker Daltonics) mass spectrometer with a Kinetex C18 column (Phenomenex). Samples were run using a linear gradient of mobile phase A (water 0.1% formic acid (v/v)) and phase B (acetonitrile 0.1% formic acid (v/v)). A representative linear gradient consisted of 0-0.5 min isocratic at 5% B, 0.5-8.5 min 100% B, 8.5-11 min isocratic at 100% B, 11-11.5 min 5% B, and 11.5-12 min 5% B. Data were collected in positive ion mode using data-dependent acquisition. All solvents used were LC-MS grade.

#### Molecular networking

Molecular networking and spectral library searching were performed using the GNPS platform as described above. Settings included a precursor mass tolerance of 0.02 *m*/*z*, fragment mass tolerance of 0.02 *m*/*z*, minimum cosine similarity of 0.6, and minimum 5 matched peaks. Data visualization was performed using the Metabolomics USI interface^50^ and spectrum–spectrum matches were evaluated manually to interpret the azithromycin suspect.

### Evaluation of flavonoid suspects

#### Mass spectrometry analysis

Untargeted metabolomics data for medicinal plants listed in the Korean Pharmacopeia were used to investigate flavonoid suspects (MassIVE dataset identifier MSV000086161). Samples were extracted using methanol. Untargeted metabolomics was performed using an Acquity liquid chromatography system coupled to a Xevo G2 Q-TOF (Waters) mass spectrometer with a BEH C18 column at 40°C (Waters Corp.; 50 mm; 2.1 mm; 1.7 μm particle size). Water (solvent A) and acetonitrile (solvent B) were used as mobile phase, both with 0.1% formic acid, and a method of 20 min (linear gradient), flow 0.3 mL/min was performed using the following settings: 0-14 min. from 5 to 95% B; 14-17 min, 95% B; 17-17.1 min from 95% to 5% B; 17.1-20 min, 5% B for equilibration of the column for the next sample. Data were collected in positive and negative ion modes using data-dependent acquisition. All solvents used were LC-MS grade.

#### Molecular networking

Molecular networking and spectral library searching were performed in the GNPS platform as described above. Settings included a precursor ion mass tolerance of 2.0 *m/z*, fragment ion mass tolerance of 0.5 *m/z*, minimum cosine similarity of 0.7, and minimum 6 matched peaks. Data visualization was performed using the Metabolomics USI interface^50^ and spectrum–spectrum matches were evaluated manually to interpret the flavonoid suspects.

### Home environment personal care products

#### Mass spectrometry analysis

The presence of polymeric suspects was investigated in the context of the HOMEChem project, a study of the indoor chemical environment (MassIVE dataset identifier MSV000083320).^31^ For full details on the experimental set-up, see Aksenov et al. (2021).^31^ Briefly, scripted activities, including cleaning and cooking, were performed in a controlled home environment. Sample collection consisted of swabbing different locations in the test house. Untargeted metabolomics was performed using a Vanquish liquid chromatography system (Thermo Scientific) coupled to a QExactive Orbitrap (Thermo Scientific) mass spectrometer with a Kinetex C18 column (Phenomenex). The mobile phase used was water (phase A) and acetonitrile (phase B), both containing 0.1% formic acid (Fisher Scientific, Optima LC/MS), employing the following gradient: 0-1 min 5% B, 1-8 min 100% B, 8-10.9 min 100% B, 10.9-11 min 5% A, 11-12 min 5% B. Data were collected in positive ion mode using data-dependent acquisition. All solvents used were LC-MS grade.

#### Molecular networking

Molecular networking and spectral library searching were performed using the GNPS platform as described above. Settings included a precursor mass tolerance of 0.02 *m*/*z*, fragment mass tolerance of 0.02 *m*/*z*, minimum cosine similarity of 0.7, and minimum 6 matched peaks.

### Alzheimer’s disease acylcarnitine analysis

#### Mass spectrometry analysis

The presence of acylcarnitine suspects was investigated in the context of the Religious Orders Study/Memory and Aging Project (ROSMAP) to study Alzheimer’s disease (MassIVE dataset identifier MSV000086415).^32^ Untargeted metabolomics was performed on human brain samples from 514 individuals with and without Alzheimer’s disease (360 Alzheimer’s disease patients, 154 healthy subjects). Human brain tissue samples were placed into tubes with 800 µl of a 1:1 mixture of H_2_O (Optima LC-MS grade W64) and MeOH (100%) containing 1 µM of sulfamethazine. The samples were homogenized using a Qiagen TissueLyser II at 25 Hz for 5 minutes, then centrifuged at 14,000 relative centrifugal force for 5 minutes before being incubated for a period of 30 minutes at -20 °C. A 200 µl aliquot of supernatant from each sample was transferred into a 96-well plate and vacuum concentrated to dryness via centrifugal lyophilization (Labconco Centrivap). Once dried, the samples were stored at -80 °C until LC-MS was performed. Untargeted metabolomics was performed using a Vanquish liquid chromatography system (Thermo Scientific) coupled to a QExactive (Thermo Scientific) mass spectrometer with a C18 column (Phenomenex Kinetex 1.7 µm C18 100 Å LC Column 50 × 2.1 mm). The mobile phase used was LC-MS grade water (phase A) and LC-MS grade acetonitrile (phase B), both containing 0.1% formic acid (Fisher Scientific, Optima LC-MS), with a flow rate set to 0.5 mL/min. Samples were injected at 95%A:5%B, which was held for 1 minute, before ramping up to 100%B over 7 minutes, which was held for 0.5 minutes before returning to starting conditions. Data were collected in positive ion mode using data-dependent acquisition to acquire MS full scan spectra, followed by MS/MS spectra of the top 5 most abundant ions. Precursor ions were fragmented once before being added to an exclusion list for 30 seconds.

#### Data analysis

Spectral library searching was performed using the GNPS platform as described above using the default GNPS spectral libraries only and including the nearest neighbor suspect spectral library. Settings included a precursor mass tolerance of 2.0 *m*/*z*, fragment mass tolerance of 0.5 *m*/*z*, minimum cosine similarity of 0.8, and minimum 6 matched peaks. Raw MS data visualization was performed using the GNPS Dashboard.^51^ Spectrum annotations corresponding to carnitines were extracted by filtering on “carnitine” in the compound name. Different spectrum annotations with near-identical precursor *m*/*z* (precursor *m*/*z* tolerance 100 ppm) and retention time (retention time tolerance 20 seconds) were merged. Feature abundances were obtained by computing extracted ion chromatograms (XICs) with *m*/*z* tolerance 100 ppm and retention time tolerance 20 seconds for all uniquely annotated acylcarnitines across all 514 raw files. Next, the Spearman correlations between all acylcarnitine XICs and the subjects’ CERAD scores (a measure of Alzheimer’s disease progression, with 1 indicating “definite” Alzheimer’s disease and 4 indicating “no” Alzheimer’s disease) were calculated and the correlation coefficients and associated p-values were recorded. Multiple testing correction of the p-values was performed using the Benjamini-Hochberg procedure, and acylcarnitines with a corrected p-value below 0.05 were considered to be significantly associated with Alzheimer’s disease. For visualization purposes the four-scale CERAD score was binarized by considering a CERAD score of 1 or 2 to correspond to positive Alzheimer’s disease patients, and a CERAD score of 3 or 4 to correspond to healthy individuals.

## Data availability

All of the data involved in this work are publicly available through GNPS/MassIVE:

## GNPS living data molecular networking

- <u>GNPS living data</u> (version November 17, 2020)
- Living data global molecular network

## Spectrum annotation using the nearest neighbor suspect spectral library

- Spectral library searching using the default GNPS libraries only: <u>part 1</u>, <u>part 2</u>, <u>part 3</u>, <u>part 4</u>, <u>part 5</u>, <u>part 6</u>, <u>part 7</u>, part 8
- Spectral library searching using the default GNPS spectral libraries and the nearest neighbor suspect spectral library: part 1, <u>part 2</u>, <u>part 3</u>, <u>part 4</u>, <u>part 5</u>, <u>part 6</u>, <u>part 7</u>, <u>part 8</u>

## Evaluation of suspect use cases

- Molecular networking of apratoxin suspects
- Molecular networking of azithromycin suspects
- Molecular networking of flavonoid suspects
- Molecular networking of home environment personal care products
- Spectral library searching of Alzheimer’s disease data
  - Using the default GNPS spectral libraries only
  - Using the default GNPS spectral libraries and the nearest neighbor suspect spectral library

Additionally, all relevant data files have been deposited to a permanent Zenodo archive at https://doi.org/10.5281/zenodo.8282733.

Metabolomics and clinical data for the ROSMAP clinic cohorts are also available via the AD Knowledge Portal (https://adknowledgeportal.org) and through request to Dr. David Bennett at Rush who provided brain samples used for analysis. The AD Knowledge Portal is a platform for accessing data, analyses, and tools generated by the Accelerating Medicines Partnership (AMP-AD) Target Discovery Program and other National Institute on Aging (NIA)-supported programs to enable open-science practices and accelerate translational learning. The data, analyses, and tools are shared early in the research cycle without a publication embargo on secondary use. Data is available for general research use according to the following requirements for data access and data attribution

(https://adknowledgeportal.org/DataAccess/Instructions). For access to content described in this manuscript see: https://doi.org/10.7303/syn30255033.1.

The nearest neighbor suspect spectral library is freely available under the CC0 license at https://gnps.ucsd.edu/ProteoSAFe/gnpslibrary.jsp?library=GNPS-SUSPECTLIST and archived on Zenodo at https://doi.org/10.5281/zenodo.8282733. Additionally, it can be used for any data analysis task on GNPS by selecting it from the CCMS_SpectralLibraries > GNPS_Propogated_Libraries > GNPS-SUSPECTLIST > GNPS-SUSPECTLIST.mgf path in the GNPS file selector dialog. Step-by-step instructions are also provided on GitHub at https://github.com/bittremieux/gnps_suspect_library and on the <u>GNPS Documentation website</u>.

Individual spectra are accessible by their Universal Spectrum Identifiers (USIs).^50,52^ The spectra displayed in **Figure 3**, **Figure 4**, **Supplementary Figure 5**, and **Supplementary Figure 6** are:

- Hexanoylcarnitine, C6:0: mzspec:GNPS:GNPS-LIBRARY:accession:CCMSLIB00003135669
- Hexenoylcarnitine, C6:1: mzspec:MSV000085561:011c:scan:2864
- Benzoylcarnitine: mzspec:MSV000085561:010c:scan:2829
- Dodecanedioylcarnitine, C12-DC: mzspec:MSV000082650:M031_48:scan:1501
- 3-Hydroxybutyrylcarnitine reference: mzspec:GNPS:TASK-015e9e338c5649a7af6715af2be98e2f-spectra/specs_ms.mgf:sca n:1
- 3-Hydroxybutyrylcarnitine suspect: mzspec:MSV000082049:20_51:scan:106
- 3-Hydroxyhexanoylcarnitine reference: mzspec:GNPS:TASK-015e9e338c5649a7af6715af2be98e2f-spectra/specs_ms.mgf:sca n:4
- 3-Hydroxyhexanoylcarnitine suspect: mzspec:MSV000085561:018b:scan:2609
- Apratoxin A: mzspec:GNPS:GNPS-LIBRARY:accession:CCMSLIB00000424840
- Apratoxin D: mzspec:GNPS:GNPS-LIBRARY:accession:CCMSLIB00000424841
- Apratoxin F: mzspec:GNPS:GNPS-LIBRARY:accession:CCMSLIB00000070287
- Apratoxin C: mzspec:MSV000086109:BF9_BF9_02_57124.mzML:scan:722
- Apratoxin A – 30.010 Da: mzspec:MSV000086109:BD5_dil2x_BD5_01_57213:scan:760
- Apratoxin F – 30.010 Da: mzspec:MSV000086109:BC11_dil2x_BC11_02_57176:scan:736
- Apratoxin A – 26.015 Da: mzspec:MSV000086109:BD5_dil2x_BD5_01_57213:scan:614
- Apratoxin A – 14.016 Da: mzspec:MSV000086109:BD11_BD11_02_57022:scan:591
- Azithromycin: mzspec:GNPS:GNPS-LIBRARY:accession:CCMSLIB00005434451
- 3’-O(desmethyl)azithromycin: mzspec:MSV000084132:Pos_C18_Aq7:scan:977
- Apigenin-8-*C*-hexosylhexoside: mzspec:GNPS:GNPS-LIBRARY:accession:CCMSLIB00004698180
- 7-*O*-methylapigenin-6-*C*-hexoside + 132.042 Da: mzspec:GNPS:TASK-38a1bd60bd094c8a97cf49d822e7f853-spectra/specs_ms.mgf:sca n:1573560
- Apigenin-8-*C*-hexosylhexoside – 30.010 Da: mzspec:GNPS:TASK-38a1bd60bd094c8a97cf49d822e7f853-spectra/specs_ms.mgf:sca n:1559636
- Apigenin-8-*C*-hexosylhexoside – 31.991 Da: mzspec:GNPS:TASK-38a1bd60bd094c8a97cf49d822e7f853-spectra/specs_ms.mgf:sca n:1559563
- Apigenin-8-*C*-hexosylhexoside – 46.005 Da: mzspec:GNPS:TASK-38a1bd60bd094c8a97cf49d822e7f853-spectra/specs_ms.mgf:sca n:1543689

## Code availability

Code to extract spectra from the molecular networks and compile the nearest neighbor suspect spectral library, as well as code notebooks to generate the figures and analyses presented in this manuscript are freely available on GitHub at https://github.com/bittremieux/gnps_suspect_library under the open source BSD-3-Clause license. A permanent code archive is available on Zenodo at https://doi.org/10.5281/zenodo.6459282.

All code was implemented in Python 3.8, and uses NumPy (version 1.19.2),^53^ SciPy (version 1.5.2),^54^ Pandas (version 1.1.3),^55^ and statsmodels (version 0.13.1)^56^ for scientific data processing, Pyteomics (version 4.4.0)^57^ to interface the UNIMOD repository,^21^ and matplotlib (version 3.5.1),^58^ Seaborn (version 0.11.0),^59^ spectrum_utils (version 0.3.4),^60,61^ Jupyter notebooks,^62^ and Cytoscape^63^ for visualization purposes.

## Supporting Information

**Supplementary Figure 1.**
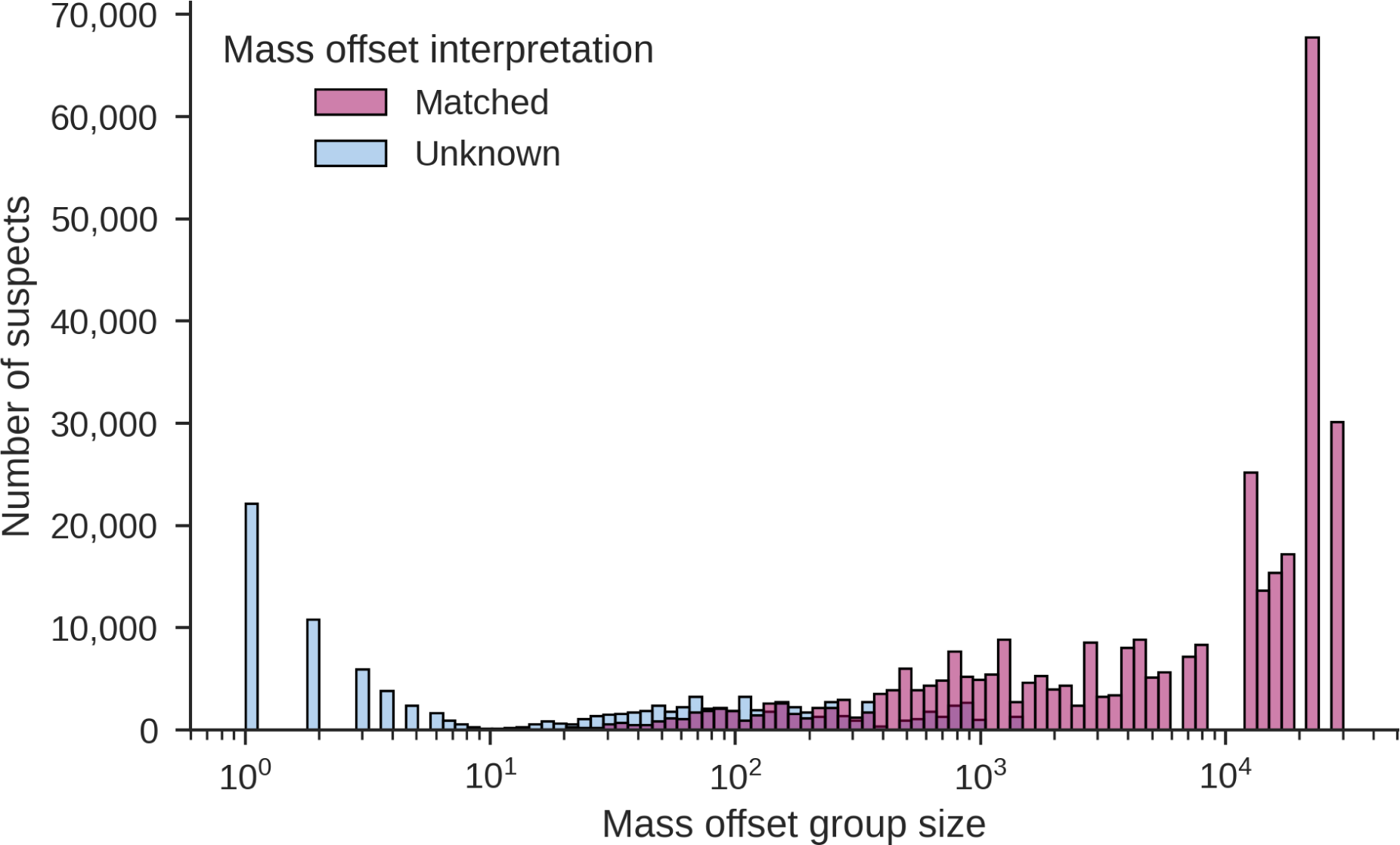
Frequency of the observed mass offsets. Several mass offsets occur hundreds to thousands of times, whereas less frequent mass offsets occur only a handful of times. Spectra with delta masses that occur fewer than ten times were not included in the final suspect library. These mass offsets could not be interpreted by matching against modifications in the UNIMOD database^21^ and a community curated list of delta masses, and are considered to be non-reproducible mass differences that likely do not correspond to real modifications.

**Supplementary Figure 2.**
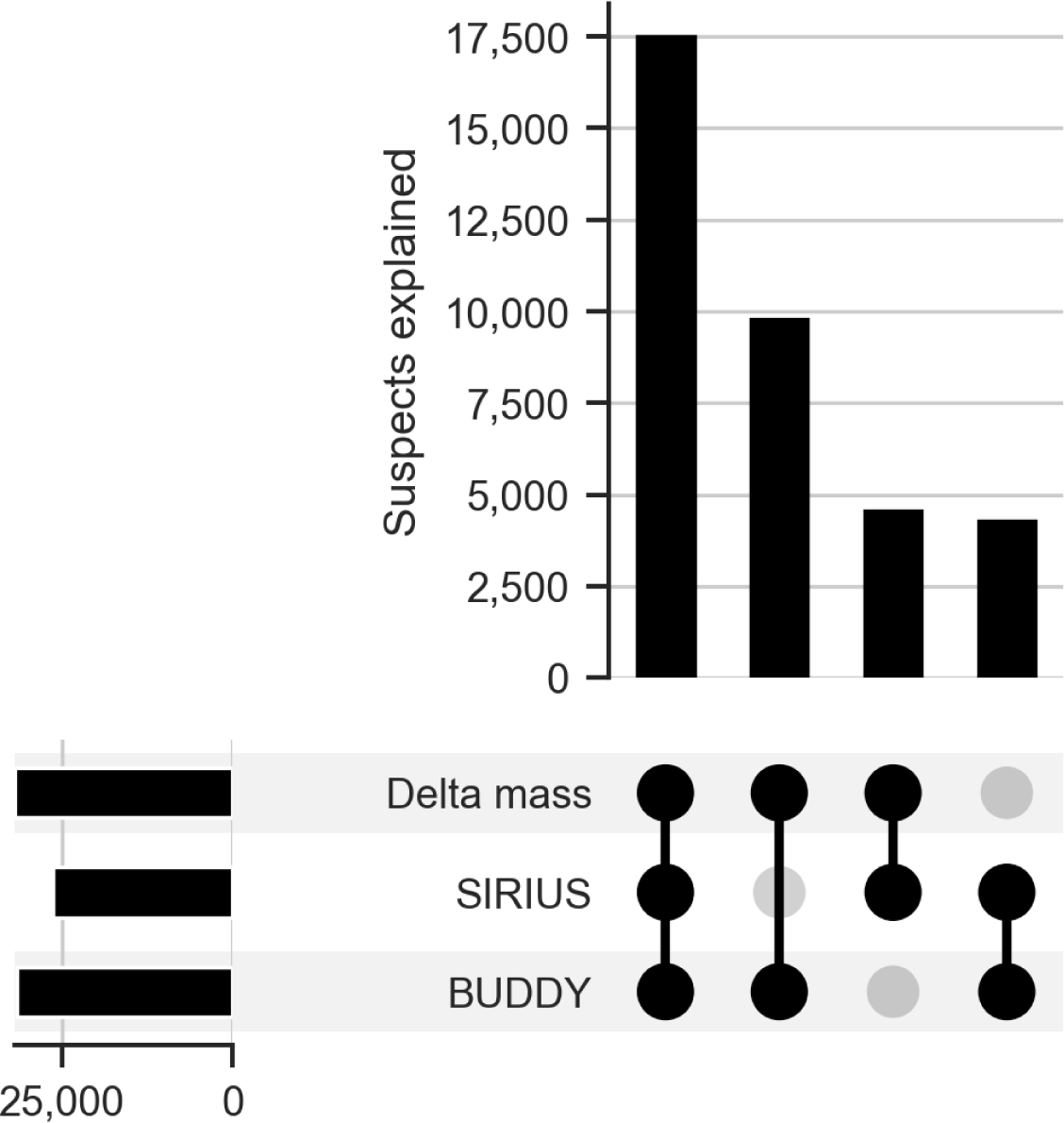
Agreement between the delta mass explanations, molecular formulas predicted by SIRIUS, and molecular formulas predicted by BUDDY for suspects for which the molecular formula of the initial molecule is known, that have a valid delta mass explanation, and for which a molecular formula could be predicted by at least SIRIUS or BUDDY. There is a large agreement between the delta mass explanations, SIRIUS, and BUDDY, with only a handful of delta mass explanations that conflict with both the SIRIUS and BUDDY predicted molecular formulas. This indicates that the delta mass explanations and the predicted molecular formulas provide complementary information that can be used to interpret the nearest neighbor suspects.

**Supplementary Figure 3.**
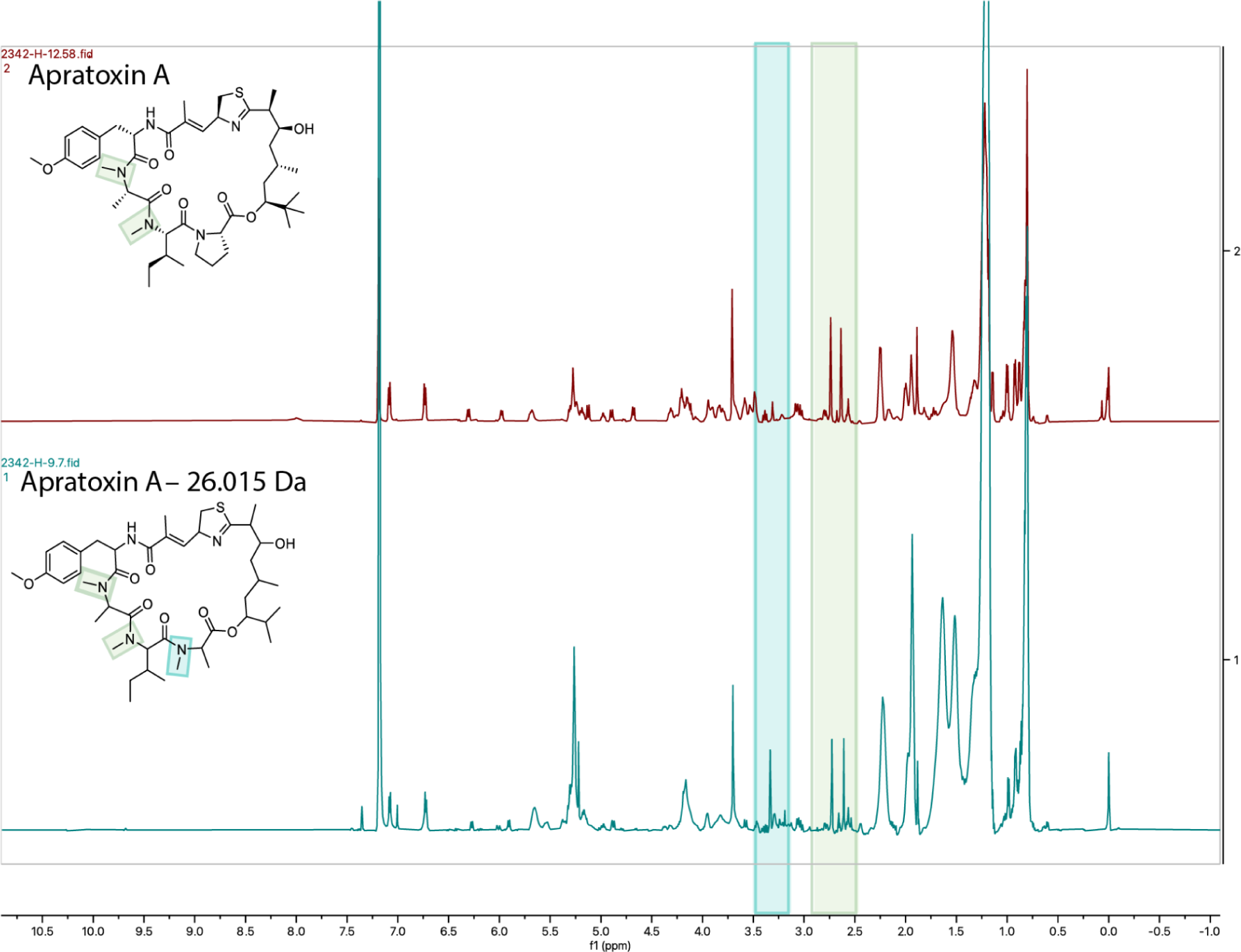
Comparison of ^1^H NMR spectra (600 MHz, CDCl_3_) of apratoxin A (top) and its related suspect (apratoxin A - 26.015 Da; bottom). Indicated by green shading are the proton signals for the *N*-methyl groups on the *N*-methyl-isoleucine and adjacent *N*-methyl-alanine at 2.71 ppm and 2.81 ppm, respectively. In the suspect there is an additional singlet proton signal observed at 3.41 ppm corresponding to the *N*-methyl-alanine adjacent to the ester bond (turquoise shading). Although the NMR results are consistent with the proposed suspect structure based on the MS/MS data, a full structure assignment was not possible due to the limited and semi-pure sample available for the NMR analysis.

**Supplementary Figure 4.**
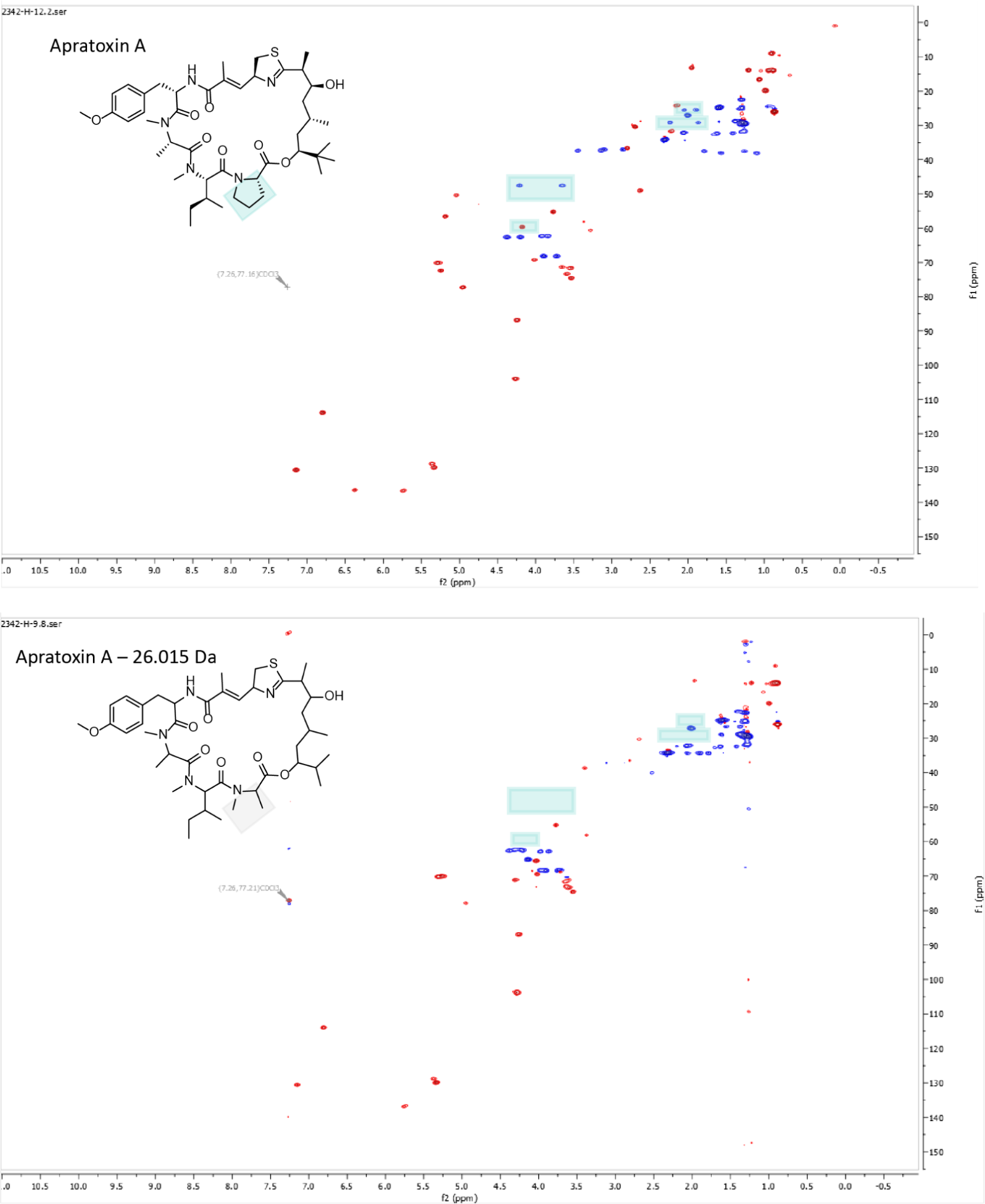
Comparison of ^1^H-^13^C HSQC spectra (600 MHz, CDCl_3_) of apratoxin A (top) and its related suspect (apratoxin A - 26.015 Da; bottom). The ^1^H-^13^C correlations associated with the proline ring (turquoise boxes) are notably absent in the suspect. Based on the MS/MS fragmentation pattern, the suspect also possesses one less methyl group in the polyketide portion of the molecule: this is possibly explained by an isopropyl rather than a *tert*-butyl group at the initiating terminus, as seen in apratoxin C. Although the NMR results are consistent with the proposed suspect structure based on the MS/MS data, a full structure assignment was not possible due to the limited and semi-pure sample available for the NMR analysis.

**Supplementary Figure 5.**
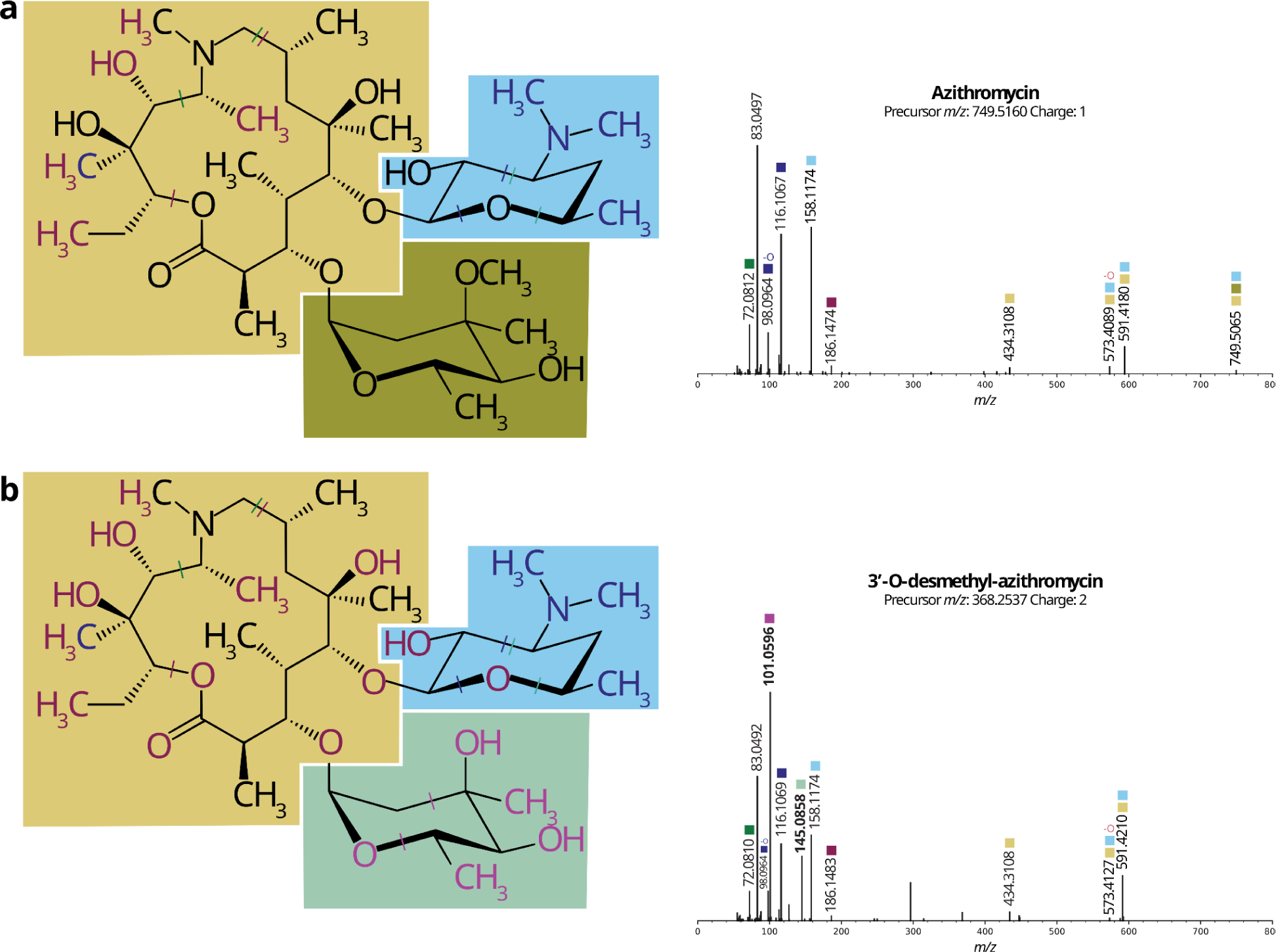
Although human breast milk is the gold standard of infant nutrition, the presence of exogenous metabolites—such as food and drugs consumed by the mother—therein is not well understood. This is especially pressing in the case of antibiotics in breast milk, as it is known that antibiotic administration in infancy can cause lasting changes in microbial colonization and host health.^64^ A public human breast milk dataset was searched for suspects related to the antibiotic azithromycin (**a**) and found specific azithromycin metabolites, including 3’-*O*-desmethyl-azithromycin (**b**), an azithromycin metabolite previously identified only in snakes.^65^

**Supplementary Figure 6.**
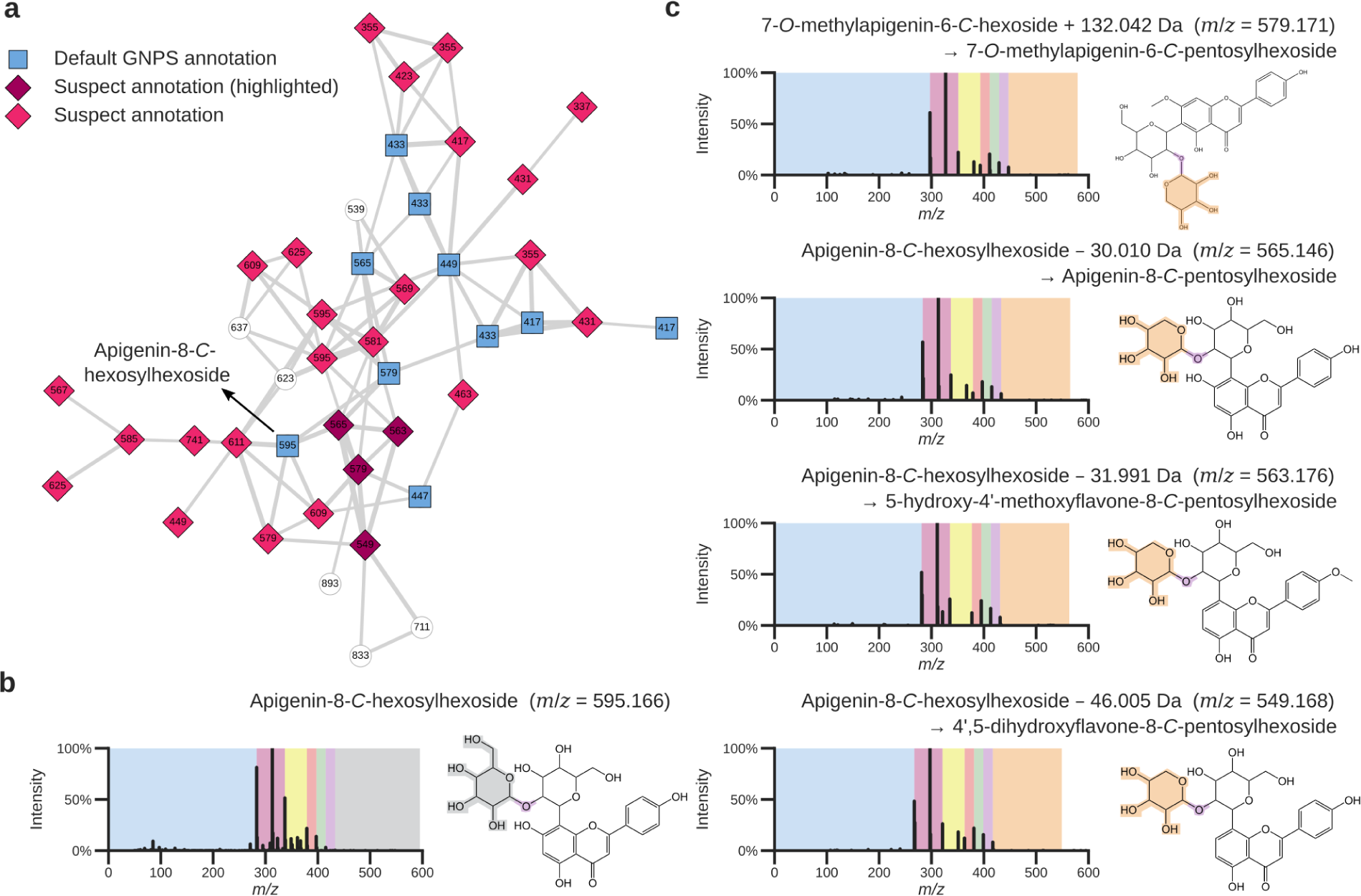
Investigation of suspects from a dataset of medicinal plants listed in the Korean Pharmacopeia.^30^ **a.** Flavonoids cluster in a molecular network created from the Korean Pharmacopeia medicinal plants dataset. The reference library hits are shown by the blue squares. The purple and pink diamonds are nodes that represent matches to the nearest neighbor suspect spectral library, with the purple diamonds matching the MS/MS spectra shown in panel c for which structures could be proposed. The white nodes are additional MS/MS spectra within the flavonoids molecular family that could not be annotated, even when utilizing the suspect library. **b.** Reference library annotation of an MS/MS spectrum matching to apigenin-8-*C*-hexosylhexoside. **c.** MS/MS spectra and structural hypotheses of apigenin-8-*C*-hexosylhexoside suspects.

**Supplementary Figure 7.**
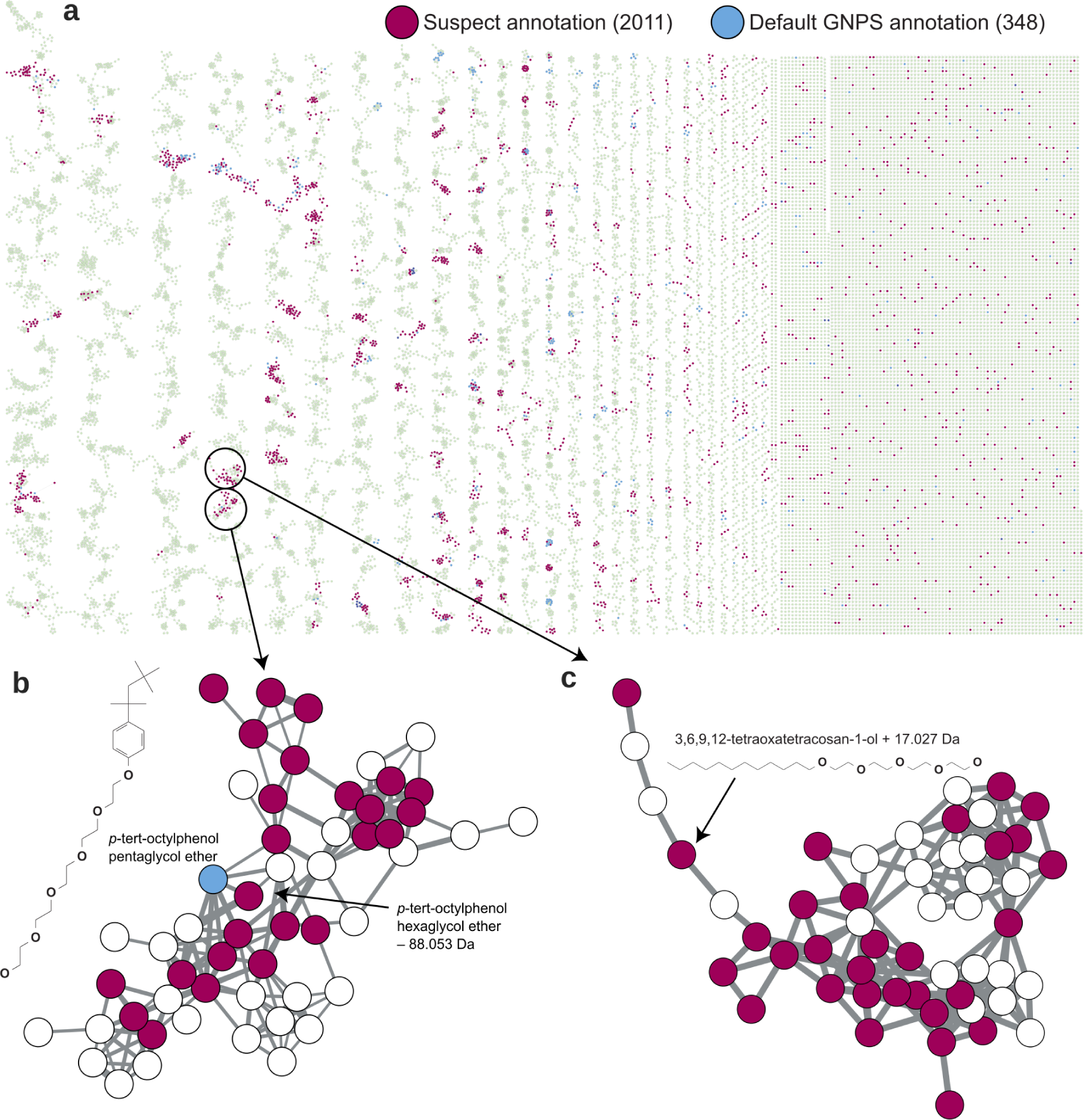
Molecular networking of the HOMEChem study to explore the chemistry of a house and how it relates to human activities within.^31^ **a.** Inclusion of the suspect library revealed a large portion of the otherwise hidden chemistry, including multiple newly annotated clusters that were found to originate from various skincare-related chemistries, in particular polyether variants. As MS/MS libraries are far from comprehensive, they contain spectra for only a small subset of possible variants of these molecules. This is especially problematic for molecules such as polyethers, as the likelihood of encountering any one particular isomer of many possible variants of polyethers, and related molecules, is very low. **b.** Example of a cluster in the molecular network where multiple spectra could be interpreted based on suspect annotations, while only a single spectrum could be annotated with conventional libraries. **c.** In the majority of cases no annotations were possible at all for skincare ingredient molecules. In contrast, using the suspect library these molecules could be readily identified. All annotations in the cluster are concordant with each other, reinforcing the suspect annotations.

